# The myocyte Nfe2l1-ubiquitin-proteasome system controls muscle fiber type and obesity-induced insulin resistance

**DOI:** 10.1101/2023.04.20.537611

**Authors:** Imke L. Lemmer, Daniel T. Haas, Fernando Rivera, Stefan Kotschi, Leonardo Matta, Nienke Willemsen, Carolin Jethwa, Joel Guerra, Alba Mena Gómez, Irmak Toksöz, Ejona Gjika, Sajjad Khani, Maria Rohm, Nick Diercksen, Phong B.H. Nguyen, Michael P. Menden, Desalegn T. Egu, Jens Waschke, Steen Larsen, Tao Ma, Zachary Gerhart-Hines, Stephan Herzig, Kenneth Dyar, Natalie Krahmer, Alexander Bartelt

## Abstract

Muscle function is an important denominator of energy metabolism and metabolic health. Adapting the myocyte proteome to energetic challenges, in response to diet or fasting, is facilitated by programs of proteostasis, but the adaptive role of the ubiquitin-proteasome system (UPS) in muscle remains unclear. Here, we show that myocyte Nuclear factor erythroid derived 2,-like 1 (Nfe2l1, also known as Nrf1) is a key regulator of skeletal muscle proteostasis and function. In mice and humans, Nfe2l1 is highly expressed in skeletal myocytes, and its loss diminishes proteasomal activity and leads to hyperubiquitylation. Mice lacking myocyte Nfe2l1 display muscle fiber type switching and insulin resistance when fed a high-fed diet. Nfe2l1 protects myocytes from ferroptosis, which is enhanced in the presence of excess lipids. In conclusion, we define a new adaptive role for the Nfe2l1-ubiquitin proteasome system in the control of skeletal muscle function and energy metabolism.

## Introduction

Skeletal muscle is a key regulator of systemic metabolism and energy homeostasis^1^. Muscle acts as a sink for glucose in an insulin-dependent manner, is a store of amino acids, and markedly contributes to whole-body energy expenditure^2^. These processes are highly adaptive and subject to environmental stimuli, nutrient availability, and physiological stress. This is particularly relevant in the context of metabolic diseases, like obesity, represented by chronic overnutrition leading to excessive weight gain and deterioration of cellular quality control mechanisms^3–5^. In skeletal muscle this metabolic imbalance is associated with compromised muscle function, fatigue, inflammation, and insulin resistance^2^. Therefore, skeletal myocytes require tightly regulated metabolic homeostasis, which on a molecular level is facilitated by the endoplasmic reticulum (ER). The ER is a central hub where protein, lipid, and parts of glucose metabolism are sensed and regulated in an adaptive manner to avoid cellular stress and inflammation^6–8^. These stress resistance mechanisms include activation of the integrated stress response, and a large body of evidence implicates the unfolded protein response (UPR) and autophagy as evolutionarily conserved quality control mechanisms involved in this process. However, much less is known about the role of the ubiquitin-proteasome system (UPS), and specifically ER-associated protein degradation (ERAD) as an alternative pathway by which cells mitigate proteotoxicity and maintain proteostasis. In this context, we and others have shown that UPS activity, rather than being static, is fine-tuned through multiple regulatory mechanisms^9^. An important transcriptional component of matching proteasomal activity to cellular demands is Nuclear factor erythroid 2-like 1 (Nfe2l1, also known as Nrf1 or TCF11), which belongs to the bZIP cap’N’collar family of transcription factors^10^. Nfe2l1 activity is subject to complex posttranslational regulation, as Nfe2l1 is tethered to the ER membrane, continuously undergoes translocation across the membrane, proteolytic cleavage, and deglycosylation before the remaining DNA-binding fragment becomes transcriptionally active. During cellular homeostasis, the Nfe2l1 fragment itself is ubiquitylated and degraded and thus virtually absent in cells. When proteasome function is suboptimal, Nfe2l1 escapes degradation and restores proteasomal activity, by binding to the antioxidant response element (ARE) in the promoter region of all proteasome subunit genes, thereby activating proteasome gene expression^11^. If proteasomal activity is compromised, e.g. in the presence of chemical proteasome inhibitors, Nfe2l1 increases the expression of proteasomal subunits, thereby adapting proteasomal activity to proteostatic needs^12^. While most of this mechanistic insight was defined in the context of cancer cells, more recent studies shed light on the physiological role of Nfe2l1-mediated proteasome function^13,14^. Nfe2l1 is activated by cold exposure in thermogenic adipocytes, and its proteasomal stimulation is a critical factor for increasing UPS activity. Compromised proteasome function, by either chemical inhibition or loss of Nfe2l1, is associated with ER stress and diminished non-shivering thermogenesis mediated by brown fat^10^. While the roles of the proteostatic mechanisms UPR and autophagy are rather well established^15–21^, the relevance of Nfe2l1-mediated adaptation of UPS for muscle function and metabolic health, specifically in obesity is still unknown.

## Results

### Nfe2l1 is linked to proteasome function in human skeletal muscle

To begin investigating the role of Nfe2l1 in skeletal muscle, we analyzed publicly available human datasets and found that *NFE2L1* is most highly expressed in skeletal muscle (Fig. 1A) and especially in fast twitch muscle fibers, yet exceeding values of the other NFE2L family member *NFE2L2* (Fig. 1B), an established regulator of muscle function. By studying single-nucleus RNA-seq datasets we determined *NFE2L1* is mainly expressed in mature myocytes, but virtually absent in other muscle cell types (Fig. 1C). Lastly, when analyzing Gene ontology (GO) pathways associated with human *NFE2L1*, the most prominent terms included the proteasome complex and ubiquitin-dependent catabolic processes (Fig. 1D). In summary, these data show that *NFE2L1* is enriched in human skeletal muscle and demonstrate a strong association between the transcription factor and proteasome function.

**Figure 1:**
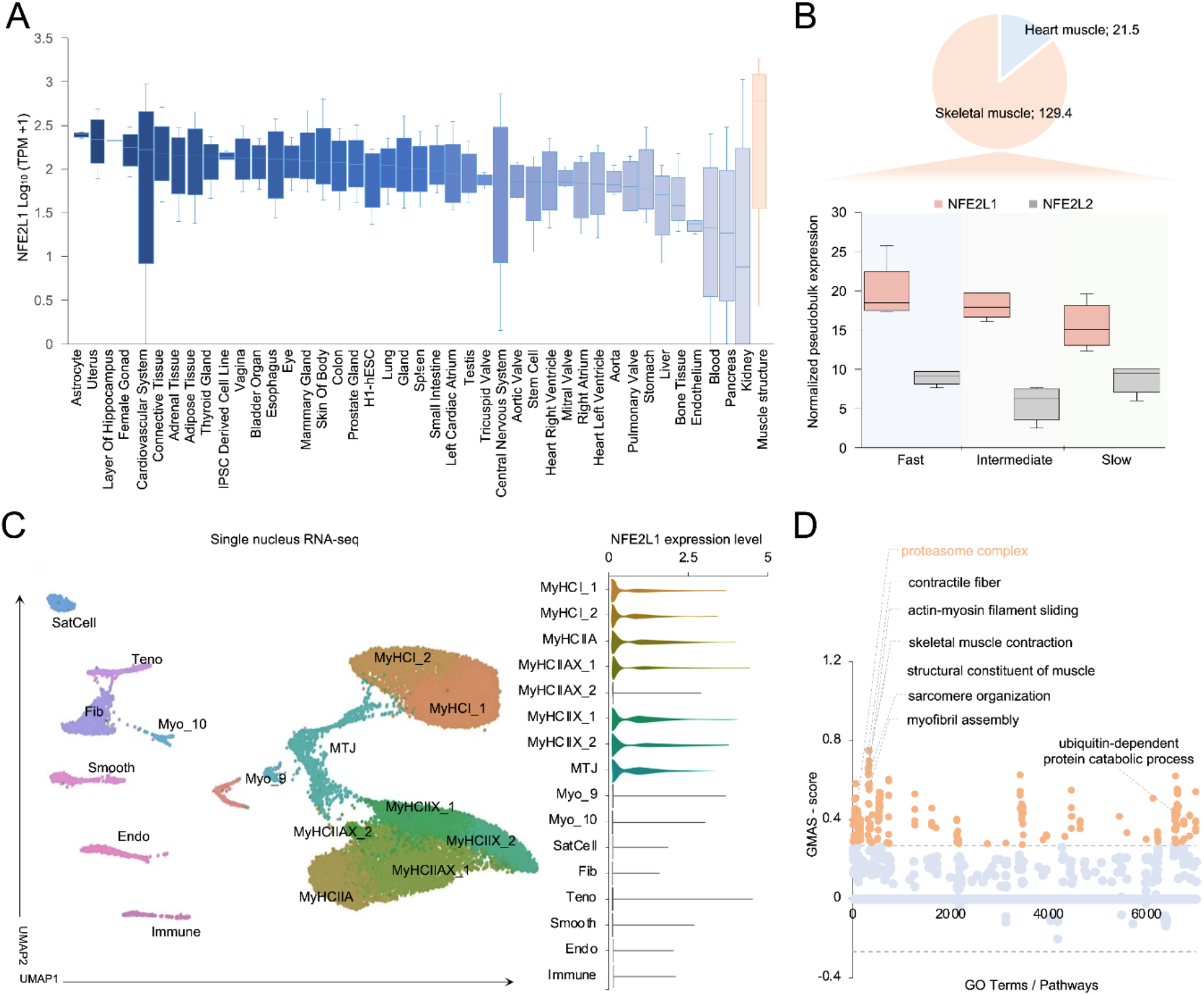
Nfe2l1 is highly expressed in human skeletal muscle and is associated with Proteasome. (A) Distribution of *NFE2L1* transcriptional expression levels (Log10 TPM + 1) across various human tissues. Skeletal muscle is highlighted on the right (orange boxes). (B) Top: Pie chart illustrating the proportional expression of *NFE2L1* in heart vs. skeletal muscle. Numerical values represent mean normalized counts per million (nCPM) in myocytes and myonuclei, respectively. Bottom: Expression levels of *NFE2L1* and *NFE2L2* across the three fiber types. Box plots represent the median and interquartile range. (C) Single-nucleus RNA-seq (snRNA-seq) analysis of skeletal muscle. Left: UMAP projection illustrating the topological clustering of distinct resident cell types and myonuclei subtypes (including endothelium, fibroblasts, immune cells, and myosin heavy chain (MyHC) phenotypes). Right: Violin plots demonstrating expression level of *NFE2L1* across each identified cell cluster. (D) GO terms and pathways associated with *NFE2L1* expression in humans. The scatter plot shows Gene-Module Association Score (GMAS ≥ 0.268, ≤ −0.268). Terms below the significance threshold are shown as light blue dots, while significant terms above the threshold are shown as orange dots.

### Myocyte-specific loss of Nfe2l1 lowers proteasome function and promotes hyperubiquitylation

To explore the biological significance of muscle Nfe2l1 we generated a transgenic Cre-loxP mouse model with a skeletal myocyte-specific deletion of *Nfe2l1* by using Acta1-Cre (*Nfe2l1* mKO). Compared to wild-type (WT) control littermates, myocyte-specific deletion of Nfe2l1 led to markedly lower proteasome subunit gene expression, depletion of NFE2L1 protein levels in muscle (Fig. 2A,B), and lower proteasome subunit protein levels (Fig. EV1A-C). Consequently, proteasomal activity in muscle tissue of *Nfe2l1* mKO animals was blunted, as assessed by fluorometric assay (Fig. 2C) and native PAGE (Fig. 2D, Fig. EV1D). Interestingly, trypsin-like activity was higher in muscle from *Nfe2l1* mKO mice, possibly indicating compensatory proteasomal activity independent of NFE2L1 and potentially facilitated by proteasome activators, which were upregulated in the mKO condition (Fig. EV1B,C). Additionally, we detected an upregulation of autophagy pathways (Fig. EV1E,F), indicating putative compensation for the loss of proteasome activity. The observation that loss of *Nfe2l1* leads to decreased levels of proteasome activity led us to investigate the impact on the proteome and the ubiquitin landscape of muscle from *Nfe2l1* mKO mice and controls by liquid chromatography paired with mass spectrometry. The impact of *Nfe2l1* mKO on the proteome was marked with 2832 differentially detected proteins in total (Fig. 2E). Moreover, the ubiquitome was profoundly altered, as seen by broad hyperubiquitylation in mKO muscles (Fig. 2F, Fig. EV1G). We quantified the amounts of different linkage-types as these determine various outcomes for the tagged proteins. Expectedly, the K48-linkage, which signals proteins for degradation via the proteasome was the most abundant and significantly higher in the *Nfe2l1* mKO muscle compared to WT controls. (Fig. EV1H). Furthermore, we observed that many proteasome subunits were ubiquitylated, suggesting that their turnover may also depend on proteasome function (Fig. EV1I). The GO analysis of the total proteome highlighted energy metabolism and filament organization as major pathways affected in muscle of mKO mice compared to control tissue (Fig. 2G). In the GO analysis of the ubiquitome, we found pathways relevant to muscle structural organization, function as well as related to the UPS (Fig. 2H).

**Figure 2:**
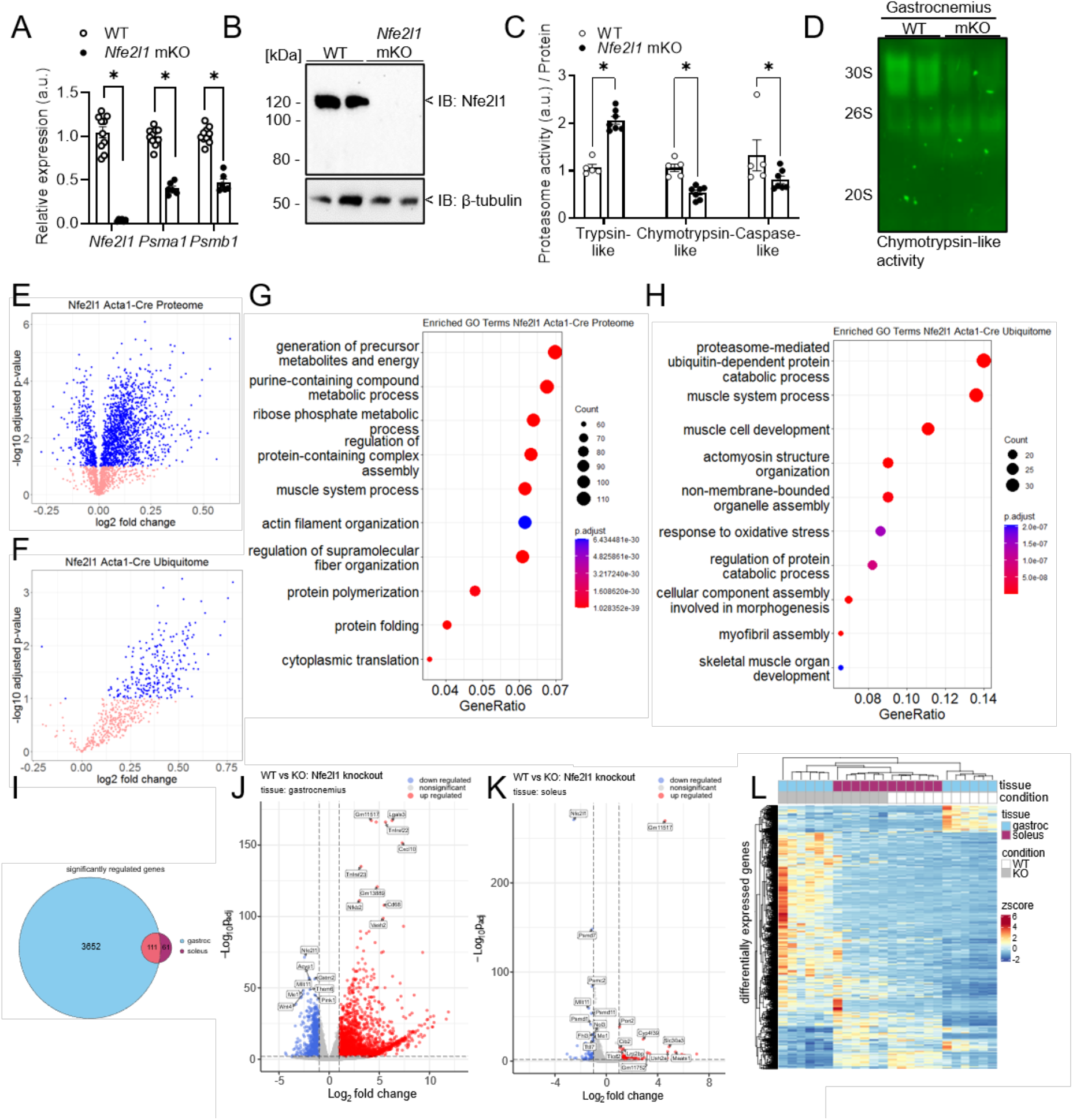
Loss of Nfe2l1 regulates proteasome function and leads to hyperubiquitylation in skeletal muscle. (A-D) Analysis of Gastrocnemius (GC) muscle from WT controls and Nfe2l1 skeletal muscle (Acta1-Cre) knock-out (mKO) mice: (A) Relative gene expression of Nfe2l1 and proteasomal subunits *Psma1* and *Psmb1* (*n* = 6 - 10 mice per group). (B) Representative Immunoblot of Nfe2l1. (C) Fluorometric assay of proteasomal activity (*n* = 5 - 7 mice per group). (D) Representative Nu-PAGE analysis of chymotrypsin-like proteasomal activity. (E-H) Assessment of GC tissue from *Nfe2l1* mKO mice and WT controls via LC-MS (*n* = 3 mice per group): (E) Volcano plot of total proteome. (F) Volcano plot of ubiquitome. (G) GO analysis of the total proteome. (H) GO term enrichment analysis of the ubiquitome. (I-K) RNAseq of GC and soleus muscle (*n* = 6 mice per group): (I) Euler diagram of significantly regulated genes in GC and soleus. Volcano plot of (J) GC and (K) soleus. (L) Heatmap of differentially expressed genes in GC and soleus of *Nfe2l1* mKO and WT mice. Data are mean ±SEM, p<0.05 by two-way analysis of variance (ANOVA) (A,C), p<0.1 by t-test (E-H) and p<0.01 by *t*-test (I-L).

### Differential impact of loss of Nfe2l1 on different fiber types

Next, we aimed to understand how loss of Nfe2l1 affects gene expression in different muscle types, and determine specific effects on individual muscle tissues, having seen a higher abundance of human NFE2L1 in the fast twitch fiber type (Fig. 1B). Therefore, we performed bulk RNA-seq of fast glycolytic gastrocnemius (GC) and slow oxidative soleus muscles from *Nfe2l1* mKO and WT mice. We confirmed the discrepancy between fast and slow fiber types we found in human muscles also in mice. Comparing the significantly regulated genes, the loss of Nfe2l1 more profoundly impacted fast GC compared to slow soleus (Fig. 2I-K). Interestingly, loss of Nfe2l1 in GC was also associated with increased immune cell and inflammation markers (Fig. 2J), suggesting increased susceptibility to damage and active repair. To further investigate these findings, we performed flowcytometry on immune cell populations in the GC and cytokines in plasma (Fig. EV2). The data show an elevated level of immune cell infiltration in GC of mKO mice compared to controls, but circulating cytokines were unchanged. Next, we compared expression patterns in soleus and GC (Fig. EV3A). We could identify differences in expression of the calcium-binding factor Cib2 and the insulin-like growth factor-binding protein Igfbp2, however, the functional significance remains to be determined. In line with our proteomics data, the GO analysis of the transcriptome (Fig. EV3B) confirmed a profound impact on genes important for UPS and energy metabolism. Additionally, it pointed towards an impact on respiration, ATP metabolic processes, and muscle cell differentiation remained unchanged. When mapping differentially expressed genes, the GC of the mKO mice clustered closer to the WT of soleus than to WT control GC itself (Fig. 2K), pointing toward a phenotypic switch of the GC tissue of mKO animals, being more similar to soleus. In summary, this unbiased global data set underscores a critical role of Nfe2l1 for UPS and proteostasis in skeletal muscle, pointing towards important roles of this process for muscle function and energy metabolism. Lastly, it shows a stronger impact on fast twitch muscle, which led us to investigate this tissue further.

### Loss of myocyte Nfe2l1 results in muscle remodeling and a lean phenotype

Our findings that UPS and specifically proteasomal activity is regulated through Nfe2l1, and the loss thereof impacts transcriptional and protein landscape, raise the question of the effect on the *in vivo* phenotype. First, we found that mKO mice displayed a lower body weight compared to controls, which was predominantly caused by lower lean mass, (Fig. 3A,B). Most prominently, GC muscle was reduced in size and weight (Fig. 3C,D), which was functionally associated with lower grip strength in *Nfe2l1* mKO mice compared to WT controls (Fig. 3E). However, expression of atrophy markers *Trim63* and *Fbxo32* were unchanged, indicating that there is no significant wasting of muscle tissue (Fig. EV3C). Histological analyses by hematoxylin and eosin staining, fiber typing, and succinate dehydrogenase staining revealed major phenotypic changes of GC (Fig. 3F). Additionally, measurement of fiber diameter showed that the more oxidative MyHCIIa fibers were larger in mKO animals (Fig. 3G), collectively indicating a shift toward a more oxidative phenotype in GC muscle. In addition, analysis of myosin heavy chain gene expression supported the phenotypic switch towards more oxidative fibers, presented by higher levels of slow-twitch *Myh7* and lower levels of fast-twitch *Myh4* expression in *Nfe2l1* mKO mice compared to controls (Fig. 3H). Furthermore, the mRNA levels of perinatal myosin heavy chains *Mhy3* and *Mhy8* were markedly higher in mKO mice compared to controls, which indicates the presence of regenerative muscle fibers (Fig. 3H). Consistent with this, the proportion of centralized nuclei was also higher in the mKO model (Fig. 3I), indicating potential regeneration and remodeling processes. Also, we observed elevated circulating creatine kinase levels in plasma (Fig. 3J) and characterizing the muscle ultrastructure with transmission electron microscopy we found that loss of Nfe2l1 led to aberrant myofibrillar organization and the mitochondria of the mKO animals seemed bulged and disrupted compared to control tissue (Fig. 3K).

**Figure 3:**
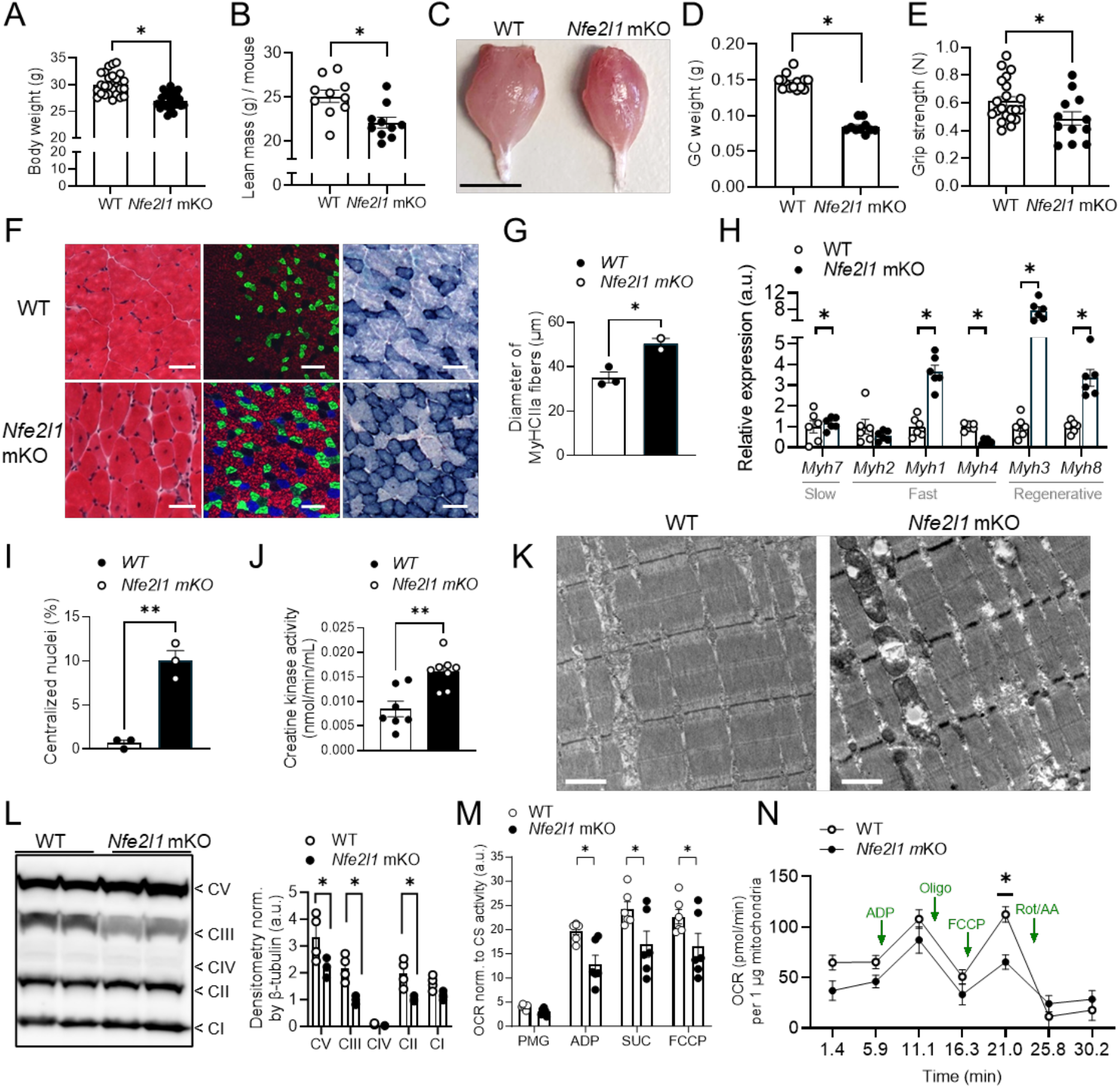
Loss of Nfe2l1 induces muscle fiber type switch and impacts mitochondrial respiration. (A) Body weight of wild-type (WT) and *Nfe2l1* mKO mice at 3 months (*n* = 26 - 27 mice per group). (B) Lean mass of mKO mice and WT control (*n* = 10 mice per group). (C) Representative photograph of WT and *Nfe2l1* mKO gastrocnemius. Scale bar: 0.5 cm. (D) Weights of gastrocnemius (GC) of WT and mKO mice (*n* = 10 - 14 mice per group). (E) Grip strength of WT controls and mKO mice (*n* = 12 - 21 mice per group). (F) Representative hematoxylin and eosin (H&E)-stained, Immunofluorescent stained fiber types MyHC-2B (red), MyHC-2A (green), MyHC-1 (blue) and succinate dehydrogenase (SDH) stained complex II in the mitochondrial respiratory chain in GC of WT and mKO mice. Scale bar: 100 µm. (G) Quantification of diameter of MyHCIIa fibers from immunofluorescent staining of GC tissue from WT and mKO mice (*n* = 2 - 3 mice per group). (H) Relative mRNA expression levels of myosin heavy chains in GC tissue from *Nfe2l1* mKO mice and WT controls (*n* = 6 mice per group). (I) Assessment of centralized nuclei in GC of WT and mKO mice (*n* = 3 mice per group). (J) Creatine kinase activity measured in plasma from *Nfe2l1* mKO animals and WT littermates (*n* = 6 - 8 mice per group). (K) Representative transmission electron microscopy (TEM) images of WT and mKO GC, longitudinal sections. Scale bar: 1 µm. (L) Immunoblot of complexes CI-CV of the electron transport chain with corresponding densitometry. (M) Oxygen consumption rates (OCR) of GC fibers from WT and mKO animals after addition of PMG (pyruvate, malate, glutamate), ADP (Adenosine diphosphate), SUC (succinate), FCCP (Carbonyl cyanide-p-trifluoromethoxyphenylhydrazone) normalized to citrate synthase (CS) activity (*n* = 6 mice per group). (N) OCR traces of isolated mitochondria from WT and *Nfe2l1* mKO mice GC tissue (*n* = 4 - 6 mice per group). Data are mean ±SEM, p<0.05 by *t*-test(A,B,D,E,G,I,J,N) or two-way analysis of variance (ANOVA) (H,L,M).

### Absence of myocyte Nfe2l1 leads to impaired mitochondrial respiration

Next, we investigated the mitochondrial phenotype and respiration further. Interestingly, GC from mKO mice contained less coenzyme Q: cytochrome-c-reductase, a component of OXPHOS complex-III (Fig. 3L). As it is established that structural alterations of mitochondria cristae impact respiration^22^, we determined mitochondrial oxygen consumption of GC muscle from *Nfe2l1* mKO and WT control mice by *ex vivo* Oroboros analysis of isolated fibers. This revealed less respiration after addition of ADP and succinate as well as reduced maximal respiration capacity in fibers of mKO mice compared to WT (Fig. 3M). To further pinpoint these results, we performed Seahorse analysis from isolated mitochondria, which confirmed the lower maximal respiration capacity in mKO GC muscle mitochondria compared to mitochondria isolated from WT controls (Fig. 3N). Notably, analysis of the metabolomic profile of GC muscle from mKO animals revealed a significant enrichment of the Warburg effect pathway (Fig. EV3D-G). Also, we find that *FGF21* and *GDF15* mRNA expression were higher in muscle of mKO mice compared to WT controls, and for GDF15 also plasma levels were elevated (Fig. EV4A-D), which could indicate an adaptive response to proteotoxic stress with potential endocrine impact on systemic energy metabolism. Moreover, loss of mitochondrial membrane potential in myocytes is associated with the induction of FGF21^23^, a myokine that is implicated in regulating energy. In summary, this set of data demonstrates a fiber-type switch from fast to slow-twitch fibers with aberrant mitochondrial bioenergetics in GC of mice lacking myocyte Nfe2l1.

### Loss of myocyte Nfe2l1 protects from weight gain due to increased energy expenditure

Next, we asked if the phenotype of the skeletal muscle *Nfe2l1* mKO mice translated into altered whole-body energy metabolism. Using indirect calorimetry, we neither found differences in respiratory exchange ratio (Fig. 4A) nor in oxygen consumption per mouse on a regular chow diet (Fig. 4B,C). However, considering the lower body weight and lean mass of mKO mice as confounding co-variates, we performed linear regression modeling of the relationship between energy expenditure and body weight and found that energy expenditure was higher in *Nfe2l1* mKO mice compared to WT controls (Fig. 4D). Interestingly, *Nfe2l1* mKO displayed higher food intake (Fig 4E), but no evident differences in physical activity or whole-body fat mass (Fig. 4F,G). As thermoregulation is a substantial component of energy expenditure in mice, we investigated whether alterations in body temperature could contribute to the elevated energy expenditure observed in *Nfe2l1* mKO animals, but this was unchanged between groups (Fig. EV4E,F). These findings raised the question how *Nfe2l1* mKO mice would respond to HFD feeding in terms of weight gain and metabolic health. We fed mKO and control mice HFD over 16 weeks and monitored body weight gain. HFD feeding led to the body weight differences becoming more pronounced, as the mKO mice were resistant to weight gain whilst the WT controls on HFD gained weight as expected (Fig. 4H,I, Fig. EV4G,H) despite food intake being higher in mKO mice compared to WT controls (Fig. EV4I). Linear regression modeling of indirect calorimetry data of energy expenditure and body weight revealed higher energy expenditure of the mKO on HFD compared to HFD-fed WT controls (Fig. 4J). While *Nfe2l1* mKO mice showed increased physical activity (Fig. EV4J), this difference alone is unlikely to account for the elevated energy expenditure, as physical activity contributes only a relatively small fraction of total energy expenditure, with the majority arising from resting metabolic and thermogenic processes^24^. Furthermore, glucose and insulin tolerance tests (GTT and ITT, respectively) (Fig. 4K,L) showed a significant difference between WT and mKO animals on HFD. The HFD-fed mKO mice displayed lower glucose and insulin tolerance compared to chow-fed mKO mice but showed a better response compared to WT animals fed with HFD. WT and mKO on chow diet showed similar GTT and ITT responses. In addition, we measured fasting plasma insulin levels and adiponectin concentrations. While there was no difference in chow-fed animals, mKO had lower insulin levels compared to WT controls when fed a HFD (Fig. 4M). Adiponectin plasma levels were lower in mKO mice on HFD compared to WT controls and interestingly also lower than *Nfe2l1* mKO on chow diet (Fig. 4N). In line with weight gain resistance of mKO animals, leptin was higher in HFD-fed WT compared to HFD-fed mKO mice (Fig. 4O), but indifferent on chow diet. Cholesterol levels increased in both genotypes on HFD yet remained significantly lower in mKO mice in comparison to WT controls (Fig. 4P). Lastly, triglyceride levels were only significantly increased in mKO mice on HFD (Fig. 4Q). To determine if the observed phenotype was influenced by developmental effects of the loss of Nfe2l1, we additionally analyzed an inducible mKO model. In this system, *Nfe2l1* was deleted only in adult animals following tamoxifen administration, thereby avoiding any risks of developmental confounders potentially associated with constitutive gene deletion. While this model was to a certain extent a phenocopy of the constitutive model, the effects were less pronounced (Fig. EV5A-I). The inducible model displayed higher insulin sensitivity than WT littermates (Fig. EV5J-O) on HFD, despite no differences in HFD-induced weight gain. In summary, these findings show that loss of myocyte Nfe2l1 is associated with relative mitochondrial dysfunction, yet increased oxidative fiber remodeling and increased energy expenditure, which protected the animals from DIO-induced detrimental metabolic effects. The differences between the constitutive and inducible model further suggest that *Nfe2l1* may play an additional role during skeletal muscle development, requiring future studies to define its specific role during this process.

**Figure 4:**
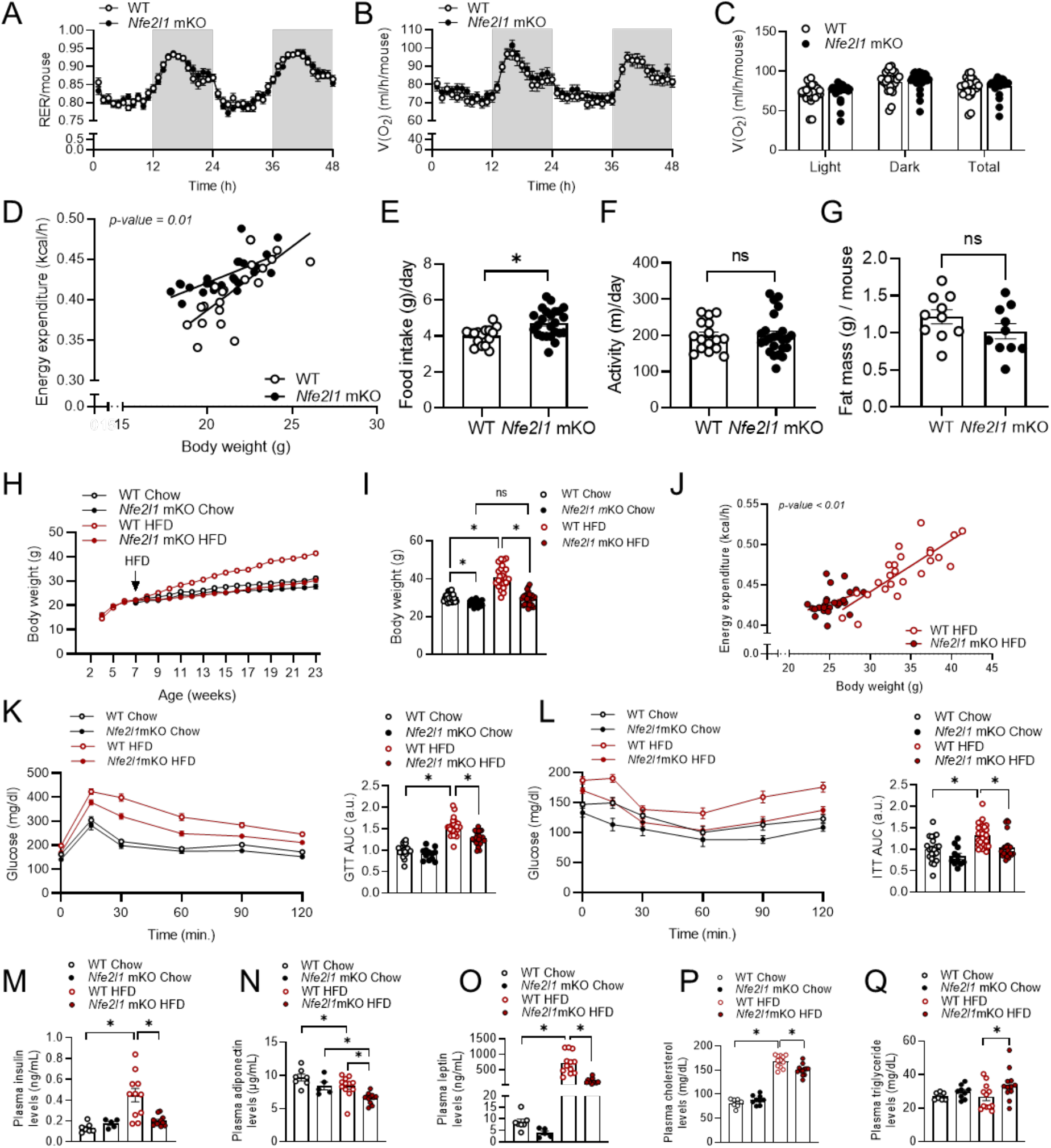
Absence of myocyte Nfe2l1 improves energy metabolism and metabolic health. (A-G) Measurements of WT and *Nfe2l1* mKO mice on chow diet at 6 weeks of age. (A) Respiratory exchange ratio (RER), (B) Traces and (C) Quantification of oxygen consumption V(O_2_) ((A-C): *n* = 26 - 28 mice per group). (D) Energy expenditure determined by indirect calorimetry over body weight (*n* = 22 - 24 mice per group). (E) Food intake. (F) Physical activity ((E,F): *n* = 16 - 24 mice per group). (G) Analysis of fat mass in 3 months old *Nfe2l1* mKO and WT mice via EchoMRI. (H) Body weight of WT controls and *Nfe2l1* mKO mice on chow or HFD over 16 weeks and (I) Body weight after 16 weeks of diet. (J) Energy expenditure after 8 weeks of HFD feeding over body weight. (H-J: (*n* = 24 - 27 mice per group)). (K) Traces and quantification (area under curve (AUC)) of intraperitoneal glucose tolerance test (GTT). (L) Traces and quantification (AUC) of intraperitoneal insulin tolerance test (ITT). (K,L: (*n* = 11 - 27 mice per group)). (M-Q) Plasma analysis of WT controls and *Nfe2l1* mKO mice on Chow or HFD after 16 weeks of diet: (M) Insulin levels, (N) Adiponectin levels, (O) Leptin levels, (P) Cholesterol levels and (Q) Triglycerides levels. ((M-Q): *n* = 5 - 12 mice per group.) The data are mean ±SEM, *P*<0.05 by *t*-test (E-G), 2-way ANOVA (C,I,K-Q) or linear modeling (D,J).

### Upstream regulators of Nfe2l1 and consequences of loss of Nfe2l1

Thus far, our results indicated that the loss of Nfe2l1 has profound impact on muscle biology, but mechanisms underlying this phenotype – while linked to UPS – remained unclear. To understand the relationship of Nfe2l1 and myocyte UPS better, we used differentiated C2C12 cells and silenced *Nfe2l1* gene expression by siRNA treatment and compared these to cells treated with control non-targeting siRNA. This strategy was successful, as both mRNA and protein levels of Nfe2l1 remained diminished even after treatment with epoxomicin, a chemical proteasome inhibitor, which stabilizes Nfe2l1 (Fig. 5A,B). Absence of Nfe2l1 led to lower levels of proteasome subunit gene expression (Fig. 5A) and, consequently lower chymotrypsin-like proteasomal activity (Fig. 5C) along with higher levels of ubiquitylated proteins after proteasome inhibition with bortezomib compared to control cells treated control siRNA (Fig. 5D). These results establish *Nfe2l1* as an adaptive regulator of proteasomal activity and ubiquitylation in cultured myocytes. Recent studies have linked proteasome function and activation of Nfe2l1 to the protection against ferroptosis, which is a lipid peroxidation, iron-dependent cell death modality^25,26^. As obesity is characterized by lipid overload and ectopic lipid deposition in muscle tissue, HFD feeding creates a lipid-rich environment prone to oxidative stress and lipid peroxidation, a hallmark of ferroptosis^27^. To mimic this ferroptotic prone state in cell culture, we treated myotubes with a mixture of ferroptosis-promoting polyunsaturated fatty acids (PUFAs)^28^, namely arachidonic acid (AA) and docosahexaenoic acid (DHA). This created a lipid-overloaded environment, which by itself enhanced oxidation of the myotubes, which was not seen by treatment with the monosaturated fatty acid (MUFA) oleic acid (OA) alone (Fig. 5E). In the PUFA-primed condition, treatment with the ferroptosis inducer RSL3 led to higher protein levels of Nfe2l1 (Fig. 5F). Correspondingly, siRNA-mediated silencing of Nfe2l1 resulted in higher oxidation in C2C12 cells than in the condition with Nfe2l1 present (Fig. 5G). In conclusion, these data show the importance of Nfe2l1 in myocytes under lipid-rich, ferroptotic conditions. Altogether, these data underline the function of Nfe2l1 in regulating muscle proteostasis, function, and the response to HFD.

**Figure 5:**
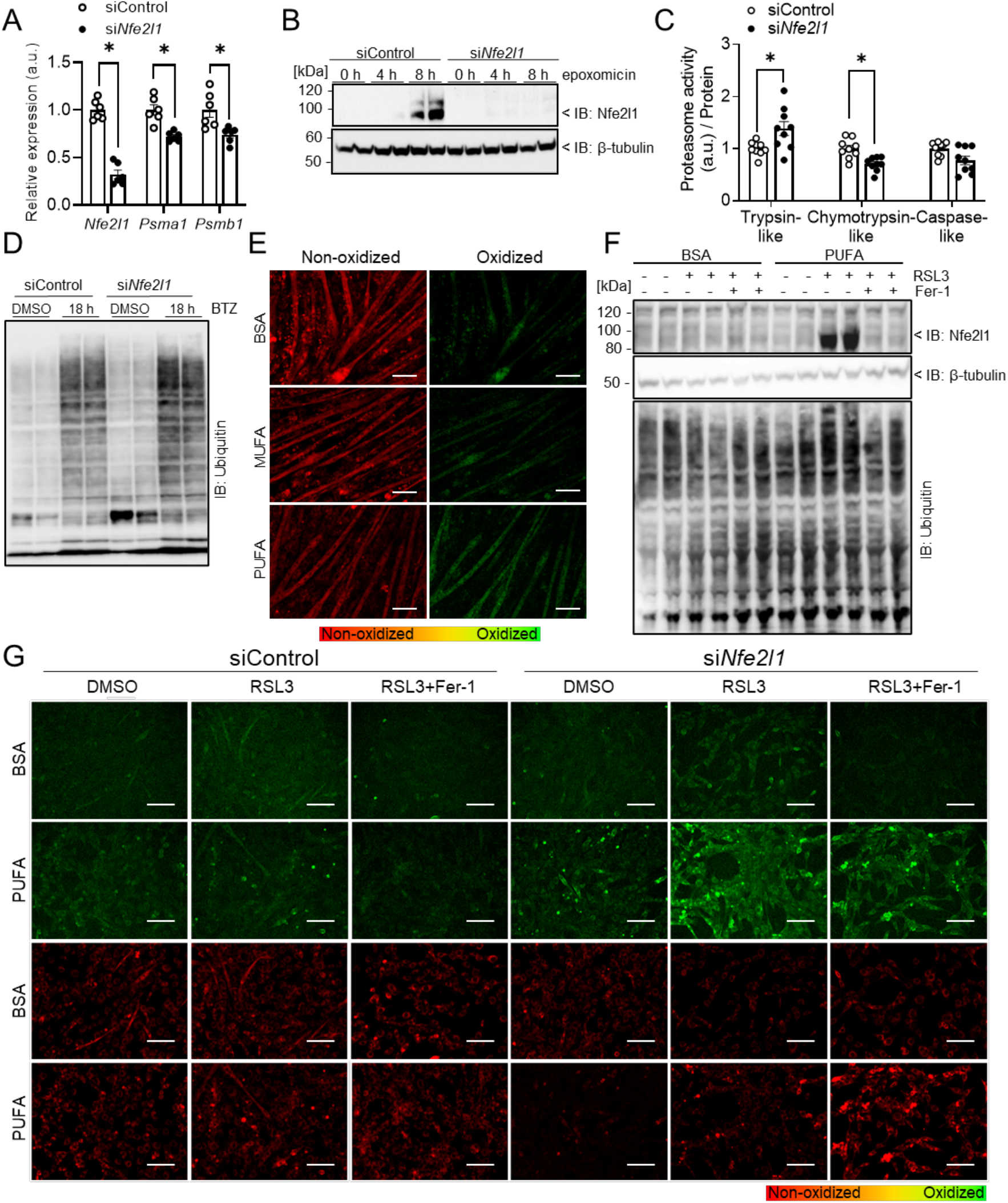
Absence of myocyte Nfe2l1 in conjunction with lipid overload increases susceptibility of cells to lipid oxidation induced by RSL3. (A-D) Protein and mRNA levels of Nfe2l1 and downstream targets in C2C12 myocytes with siControl or siNfe2l1: (A) Relative gene expression of Nfe2l1 and proteasomal subunits Psma1 and Psmb1 (*n* = 3 technical replicates from 2 independent experiments). (B) Immunoblot of Nfe2l1 with 100 nM epoxomicin treatment. (C) Fluorometric assay of proteasomal activity (*n* = 3 technical replicates from 3 independent experiments) and (D) Immunoblot of ubiquitylated proteins with bortezomib (BTZ) treatment. (E) Representative images of BODIPY 581/591 C11 stain of differentiated C2C12 myotubes treated with 0.1 mM of BSA, MUFA or PUFA for 24 h. Scale bar 100 µm. (F) Immunoblot of Nfe2l1 and ubiquitylated proteins from C2C12 myotubes pre-treated with 1 mM BSA or PUFA for 24 h, following treatment with 1 µM RSL3 and 10 µM Ferrostatin-1 (Fer-1) for 24 h. (G) Representative images of BODIPY 581/591 C11 staining in cells treated with 0.1 mM BSA or PUFA, and 0.5 µM of RSL3 and 10 µM Fer-1 for 24 h. Scale bar 100 µm. The data are mean ±SEM, *P*<0.05 by 2-way ANOVA (A,C).

## Discussion

Proteostasis is a critical component of cellular integrity and adaptation to stress. Adapting the proteome to metabolic demand, especially in tissues experiencing large fluctuations in metabolic rate such as skeletal muscle is important to prevent cellular stress and dysfunction. Here, we show that proteasomal activity and management of ubiquitin levels in muscle is a critical process regulated by myocyte Nfe2l1. Foremost, proteasomal degradation is a mechanism to dispose of and/or recycle damaged or obsolete proteins. In muscle tissue not only protein synthesis but also degradation plays a pivotal role in adaptation to different stimuli and environments^29,30^. A critical degradation pathway in the muscle is the UPS^2^ and periods of adaptation will cause more ubiquitylation, as a means for marking proteins for degradation, and thereby create higher proteasomal demand^31^. We show that Nfe2l1 is critical for myocyte proteasome function. However, loss of Nfe2l1 did not completely abolish proteasomal activity (Fig. 2C-D), which indicates that the baseline of proteasome subunit gene expression is likely mediated by a different, less adaptive mechanism, which could be mediated by e.g., FoxO1^32^.

Furthermore, in cells or tissues lacking Nfe2l1 we found that trypsin-like protease activity of the proteasome is increased (Fig. 2C and D and 5C), along with an elevated protein abundance of “Psme” subunits, also known as proteasomal activators (Fig. EV1A-C), indicating a potential compensatory posttranslational modification of proteasome function. However, overall proteasomal activity was lower (Fig. 2C and D and 5C) and ubiquitin levels higher (Fig. 2F and 5D and Fig. EV1H), demonstrating the role of Nfe2l1 determining rates of proteasome content in myocytes. Next to changes in total amount and activity, we also found that proteasome subunits accumulated in a hyperubiquitylated state (Fig. EV1I), but the relevance of this observation will need further investigation. Likely, somewhat paradoxically, this indicates that proteasome subunits themselves are regulated by proteasomal degradation and in the absence of proper proteasomal activity, these subunits might further exert dominant negative effects on proteasome assembly and/or activity. It will be interesting to understand the composition of proteasomes under these conditions, as this might reveal new biochemical levers of managing UPS activity. The accumulation of ubiquitylated proteins induces cellular stress and this is evident in our study. As polyubiquitin chains linked by K48 are the main driver for proteasomal degradation and Nfe2l1 loss impairs proteasome activity, a subsequent accumulation of K48 was anticipated (Fig. EV1H). Nevertheless, other linkages were also significantly upregulated and suggest a greater impact of the loss of Nfe2l1 than solely on the UPS. It will be relevant to investigate these sites and proteins in more detail to potentially discover new regulators of muscle biology. In terms of generalized muscle function and health, there are parallels between impaired autophagy and diminished proteasome function. When autophagy is suppressed, such as in Pompe disease, this is associated with ubiquitylated protein inclusion bodies and markers of muscle damage^33^. In general, skeletal muscle weakness is associated with the accumulation of high molecular weight proteins and ubiquitylated protein inclusions, for example in sporadic inclusion body myositis and hereditary inclusion body myopathies^34^, including those caused by mutations in valosin-containing protein VCP, also known as p97, which interestingly is a critical component of ERAD and Nfe2l1 translocation across the ER membrane^35^. There seems to be profound crosstalk between proteostatic mechanisms in muscle, as we found in the proteome of *Nfe2l1* mKO muscle that autophagy pathways are markedly upregulated, including p62 and LC3B levels (Fig. EV1E,F).

Nfe2l1 is also linked to mTORC1 signaling, as sustained genetic activation of mTORC1 promotes Nfe2l1 activation, proteasomal activity, and muscle atrophy^21^. The accumulation of ubiquitylated proteins is hallmark of muscle dysfunction and vice versa is causally linked to muscle damage when autophagy or UPS are compromised. Markers of muscle atrophy were not increased upon loss of Nfe2l1 (Fig. EV1G), indicating that the phenotype of mKO should not be classified as atrophic, but rather a lean, remodeling-driven muscle phenotype. We also observed distinct changes in GC vs soleus, with GC being more affected by Nfe2l1 loss than soleus (Fig. 2I-L), pointing to fiber type-specific demands for proteasome function and a potentially altered regenerative capacity of these fibers. Puzzling observations that require further discussion are the mitochondrial phenotype and altered energy metabolism of *Nfe2l1* mKO mice. On the one hand, there is aberrant mitochondrial function in isolated fibers and mitochondria of mKO mice compared to WT controls (Fig. 3M and N). This has been indicated in previous studies using a deletion model of the upstream regulator of Nfe2l1 leading to mitochondrial fragmentation and impaired function^36^, underlining the importance of Nfe2l1 function for mitochondrial integrity. In addition, there is a fiber-type shift in GC muscle, but not soleus, to a more oxidative, mitochondria rich phenotype (Fig. 3F to H). Regardless, on a regular diet, there is no baseline difference in whole-body RER, but increased energy expenditure and food intake. Our interpretation is that this is congruent with an energy wasting phenotype based on energy tradeoffs between mitochondrial and glycolytic metabolism and indeed, based on metabolomics, muscle of *Nfe2l1* mKO mice shows a Warburg-like metabolism (Fig. EV3F). In addition to cytosolic degradation, proteasomal protein degradation also takes place at the ERAD or in the nucleus. Clearly, disturbances in UPS and ERAD cause ER stress and inflammation^10^, which we also observe in our model (Fig. EV2). It is plausible that the inflammation contributes to energy wasting. Understanding these tradeoffs between nutrient fluxes and metabolic efficiency will require tracer studies to pinpoint the exact underlying changes and will further enhance our understanding of metabolic adaption of skeletal muscle. Uncoupling of mitochondria and loss of mitochondrial membrane potential in myocytes are associated with the induction of FGF21^23^, a myokine that is implicated in regulating energy metabolism. We find that FGF21 and GDF15 expression were higher in muscle of mKO mice compared to WT controls, and for GDF15 plasma levels were also elevated (Fig. EV4A-D), which could indicate an adaptive response to proteotoxic stress with potential endocrine impact on systemic energy metabolism. As it has been shown, that GDF15 cannot only reduce body weight by regulating appetite, but also maintain a higher energy expenditure in the muscle during food restriction, leading to an even greater weight loss. Furthermore, it has been shown to have a positive impact on glucose tolerance and insulin sensitivity^37^. Thus, these two endocrine factors could play a role in these positive effects of the loss of Nfe2l1. This is particularly interesting compared to other Nfe2l1-associated outcomes in mouse models of metabolic fatty liver disease^38^ or thermogenesis^13^, in which loss of Nfe2l1 and subsequent proteasomal impairment are clearly and exclusively detrimental. In contrast, here we show that loss of Nfe2l1 in myocytes is well tolerated and results in a favorable metabolic profile. This is in line with the notion that skeletal muscle responds to stress in a more differentiated fashion such as certain levels of reactive oxygen species in exercise are required for its beneficial adaptation, yet also containing oxidative stress through an interplay alongside other transcriptional regulators like Nfe2l12, also known as Nrf2^39–41^.

This balance appears particularly important in conditions characterized by chronic oxidative stress and metabolic imbalance. Obesity represents such a pathophysiological state, characterized by insulin resistance, aberrant lipid metabolism, and mitochondrial dysfunction, all of which lead to increased reactive oxygen species and protein damage^40^, causing loss of muscle mass and function^42^. Other studies have found Nfe2l1 to be a central regulator of cellular adaptation to oxidative and ferroptotic stress and safeguards the cells from cell death^25,26,43^, but the role in ferroptosis in the muscle is unclear. Our findings suggest that in a lipid-rich, PUFA-overloaded environment Nfe2l1 accumulates upon ferroptotic induction, indicating a comparable protective role as displayed in other studies. Consequently, the absence of Nfe2l1 seems to lower the threshold for lipid peroxidation, rendering myocytes more subjected to the detrimental effects of chronic lipid overload and redox imbalance.

In conclusion, our study demonstrates the pivotal role of Nfe2l1 for muscle proteostasis and energy metabolism. We highlight the impact of proteasome function and turn-over of ubiquitylated proteins on muscle fiber identity and mitochondrial respiration. Additionally, we identified a role of ferroptosis in the regulation of myocyte Nfe2l1. Advancing our understanding of the regulation of Nfe2l1 might generate a therapeutic handle to preserve muscle mass and healthy metabolism in obesity and beyond.

## 2 Materials and methods

### 2.1 Mice husbandry, tissue collection, and plasma analysis

All animal experiments were performed according to procedures approved by the animal welfare committees of the government of Upper Bavaria, Germany (ROB-55.2-2532.Vet_02-20-32 and ROB-55.2-2532.Vet_02-21-56) and performed in compliance with German Animal Welfare Laws. Animals for dietary induced obesity (DIO) experiments were C57BL/6JRj with 16 weeks of HFD and age-matched chow fed controls purchased from Janvier Labs. The Nfe2l1 floxed mice described previously^13^ mice were crossed with B6.Cg-Tg(ACTA1-cre)79Jme/J (The Jackson Laboratory, stock no. 006149), as well as the tamoxifen inducible Acta1-CreERT2 to create constitutive and inducible Nfe2l1 myocyte-specific knockout (mKO) mice and WT control littermates carrying floxed Nfe2l1 alleles yet being Cre negative (Cre^−^/CreERT2^−^). The CreERT2 mice were injected with 3 doses of 80 mg/kg Tamoxifen (Sigma) in Miglyol (Caelo) on days 0, 2 and 4 of 7 weeks of age to induce the Nfe2l1 mKO. The mice were bred in standard conditions (License: ROB-55.2-2532.Vet_02-30-32 and ROB-55.2-2532.Vet_02-21-56)and housed in individually ventilated cages (IVCs) at room temperature (22 °C), with a 12 h light-dark cycle, and fed chow-diet or high-fat diet (HFD) with 60 kJ % lard (Sniff, D12492) and water ad libitum. Indirect calorimetry was measured in a PromethionCore by Sable Systems. Body composition was measured by EchoMRI™. Mice were anesthetized with a xylazine/ketamine (30/150 mg/kg) (WDT) prior to blood sampling and perfusion with PBS (Thermo Fisher Scientific) with heparin (50 U/mL). Collected tissues were snap-frozen in liquid nitrogen and stored at −80 °C. Plasma Leptin (R&D Systems), Adiponectin (R&D Systems) and Insulin (Crystal Chem) were analyzed via ELISA kits according to the manufacturer’s instructions. Plasma cholesterol and triacylglycerol levels were measured using a Beckman Coulter AU480 Chemistry Analyzer (Beckman Coulter, Brea, CA, USA). Glucose and insulin tolerance tests were conducted after 6 h of fasting by intraperitoneal administration of 1 mg/g glucose or 0.75 U/kg insulin in 0.2 % wt/vol BSA in PBS. Forelimb grip strength was assessed in triplicates via the Bioseb grip strength test instrument (GS3).

### 2.2 Cell culture, treatments, and reverse transfection

For cellular experiments, we used immortalized C2C12 mouse myoblasts (Sigma-Aldrich, 91031101). All cells were kept at 37 °C, 5 % CO_2_ in DMEM Glutamax (Thermo) supplemented with 10 % v/v FBS (Sigma) and 1 % v/v Penicillin/Streptomycin (PenStrep (Sigma)). They were continuously kept between 30-80% confluency and split three times per week. Cells were used until their passage number exceeded 40. For differentiation, C2C12 cells were grown to 95 % confluency followed by differentiation with DMEM Glutamax (supplemented with 2% v/v horse serum (Sigma) and 1 % v/v PenStrep). The medium was replaced every day for 6 days. Differentiated cells were treated with DMSO (Sigma), 100 nM bortezomib (SelleckChem) or 100 nM epoxomicin (Merck) before harvest. For ferroptosis experiments C2C12 cells were pre-treated with either oleic acid (18:1) (Sigma) or with a 3:1 ratio of arachidonic acid (AA, 20:4) (Cayman Chemical) and docosahexaenoic acid (DHA, 22:6) (Cayman Chemical) conjugated to 5% BSA (Sigma), consequently treated with RSL3 (SelleckChem) and Ferrostatin-1 (SelleckChem) as indicated. For staining, cells were fixed for 15 min at room temperature with 4% paraformaldehyde (PFA) and then washed and stained with 3 µM of BODIPY 581/591 C11 (Thermo) for 30 min in the dark, followed by washing twice with PBS. Images were acquired using an EVOS5000 (Thermo). For *in vitro* gene silencing, SMARTpool small interfering RNA (siRNA, Dharmacon) in Lipofectamine RNAiMAX transfection reagent (Thermo), was used according to manufacturer’s instructions. Reverse transfection was performed on day two of differentiation using 30 nM siRNA. New differentiation medium was added 24 h after transfection and cells were treated 48 h after transfection and harvested subsequently.

### 2.3 Gene expression analysis

NucleoSpin RNA kit (Macherey Nagel) was used according to the manufacturer’s instructions, for RNA extraction. RNA concentrations were measured with a NanoDrop spectrophotometer (Implen). For complementary DNA (cDNA) preparation 500 ng RNA were added to 2 μL Maxima H Master Mix (Thermo) and the total volume adjusted 10 µL with H_2_O. The cDNA solution was diluted 1:40 in H_2_O. Relative gene expression was measured with qPCR. Per reaction, 5 µL PowerUp™ SYBR Green Master Mix (Thermo), 4 µL cDNA, and 1 µL of 5 µM primer stock (Supplementary Table 1) were added. A Quant-Studio 5 RealTime PCR system (Thermo) 2 min 50 °C, 10 min 95 °C, 40 cycles of 15 sec 95 °C, 1 min 60 °C) was used for measurement of Cycle thresholds (Cts) of genes of interest. Data was normalized to *TATA-box binding protein (Tbp)* levels by the ΔΔCt-method. Relative gene expression is displayed as fold change to the appropriate experimental control groups. RNA sequencing (RNAseq) was performed by Novogene (UK). For library sequencing Illumina NovaSeq platforms were used, applying a paired-end 150 bp sequencing strategy (short-reads).

### 2.4 Protein isolation/analysis and citrate synthase activity

Protein extraction and immunoblotting were performed as previously described^44^. Briefly, cells or tissues were lysed in RIPA buffer (50 mM Tris pH 8 (Merck),150 mM NaCl (Merck), 5 mM EDTA (Merck), 1 % v/v IGEPAL® CA-630 (Sigma), 0.5 % v/v sodium deoxycholate (Sigma), 0.1% v/v SDS (Roth)) containing cOmplete mini protease inhibitors (Sigma) and PhosSTOP^TM^ (Roche Diagnostics) using a TissueLyser II (3 min, 30 Hz; Qiagen). Lysates were centrifuged for 15 min (21,000 *g*, 4 °C) and supernatant was collected. Protein concentrations were determined using the Pierce BCA Protein Assay (Thermo) according to the manufacturer’s instructions. 15-35 µg protein per sample were denatured with 5 % v/v 2-mercaptoethanol (Sigma) in LDS sample buffer (Thermo) for 5 min at 95 °C before they were loaded in Bolt™ 4–12 % Bis-Tris (Thermo) or NuPAGE^TM^ 12% (Thermo) gels. After separation, proteins were transferred onto a 0.2 µm PVDF membrane (Bio-Rad) using the Trans-Blot® Turbo™ system (Bio-Rad) at 25 V, 1.3 A for 7 min. For determination of protein concentration, the membrane was stained in Ponceau S (Sigma). Blocking occurred in Roti-Block (Roth) at room temperature for 1 h. Primary antibodies (Supplementary Tab. 2) were incubated overnight at 4 °C. Blots were washed 3 x for 7 min with TBS-T (200 mM Tris (Merck), 1.36 mM NaCl (Merck), 0.1% Tween 20 (Sigma)) and secondary antibodies (Supplementary Tab. 2) applied for 1 h at room temperature. The membranes were washed in TBS-T and developed using SuperSignal West Pico PLUS Chemiluminescent Substrate (Thermo) in a ChemiDoc imager (Bio-Rad). We normalized the protein bands to beta-tubulin with Image Lab software (Bio-Rad). The uncropped blots can be found in Supplemental figure 1. For citrate synthase activity, 15-30 µg protein was added to the reaction buffer (20 mM Tris-HCl, pH 8.0 (Merck), 0.42 mM acetyl-coenzyme A (Sigma), 0.1 mM DTNB (Sigma)) and incubated for 5 min at 37 °C. Subsequently, 0.5 mM oxaloacetate was added, and emission measured at 412 nm over a course of 5 min.

### 2.5 Proteasome activity

Cells were lysed in lysis buffer (40 mM TRIS, pH 7.2 (Merck), 50 nM NaCl (Merck), 5 mM MgCl_2_ (hexahydrate) (Merck), 10% v/v glycerol (Sigma), 2 mM ATP (Sigma), 2 mM β-mercaptoethanol (Sigma). Proteasome Activity Fluorometric Assay II kit (UBPBio, J41110) was used according to manufacturer’s instructions for assessment of activity. Fluorescence was measured in the Plate reader (Tecan). This assay allowed for chymotrypsin-like, trypsin-like, and caspase-like activity measurement. The results were normalized to protein content using Bio-Rad Protein Assay Kit II (Bio-Rad) according to manufacturer’s instructions.

### 2.6 Native PAGE

The protocol for in-gel proteasome activity assay and following immunoblotting was previously described in detail^45^. Briefly, tissues were lysed in Lysis buffer (50 mM Tris-HCl pH 7.5, 5 mM MgCl_2_, 2 mM DTT, 10% v/v glycerol, 2 mM ATP, 0.05% v/v Digitonin), with a phosphatase inhibitor (PhosStop^TM^, Roche). The suspensions were kept on ice for 20 min and subsequently centrifuged twice. Protein concentration was determined with Bio-Rad Protein Assay Kit II and 15 μg sample protein was loaded on a NuPAGE 3-8% Tris-Acetate gel (Thermo). The gel was run for 4 h at a constant voltage of 150 V. Next, the gel was incubated in an activity buffer (50 mM Tris, 1 mM DTT, 1 mM MgCl_2_) with 0.05 mM chymotrypsin-like substrate Suc-Leu-Leu-Val-Tyr-AMC (Bachem) at 37 °C for 30 min. The fluorescent signal was measured via a ChemiDoc MP (Bio-Rad).

### 2.7 Histology and EM

For Hematoxylin and eosin staining tissues were fixed in 4% formalin (Sigma) and dehydrated overnight (70% EtOH, 96% EtOH, 100% EtOH, Xylene (Sigma)) in an ASP200S (Leica) and consequently embedded in paraffin (Roth). Sections of 4-6 µm were produced using a microtome (Leica). Deparaffination and rehydration was performed using standard procedure by incubating in Xylene (2 x 5 min), 100 % EtOH (2 × 10 dips), 96% EtOH (5 dips), 70% EtOH (3x), ddH_2_O (10x). Staining occurred by incubation in hematoxylin, water, eosin (5 min each) and subsequent dehydration. Slides were mounted with Histokitt II (Roth) and images were taken with DMi8 (Leica). For SDH staining cryosections were produced with a Cryotome CM3050S (Leica) at 4-6 µm and stained with nitroblue tetrazolium (1:1000 w/v, Thermo) diluted in SDH stock solution (0.2 M sodium succinate, 0.2 M PBS, Sigma). Sections were stained for 45 min at 37 °C, following rinsing with acetone and dehydrating with ethanol and xylene. For immunofluorescent staining sections were incubated with 4% mouse on mouse (MOM) IgG blocking solution (Vector) at room temperature, briefly washed with PBS and incubated for 1 h at 37 °C with a mix of primary antibodies (DSHB (University of Iowa)) consisting of: BA-D5 (IgG2b, supernatant, 1:100 dilution) specific for MyHC-I, SC-71 (IgG1, supernatant, 1:100 dilution) specific for MyHC-2A and BF-F3 (IgM, purified antibody, 1:100 dilution) specific for MyHC-2B. Subsequently, washed 3 times with PBS and incubated with secondary antibodies (Jackson ImmunoResearch) for 45 min at 37 °C: goat anti-mouse IgG1, conjugated with DyLight488 fluorophore, goat anti-mouse IgG2b, conjugated with DyLight405 fluorophore, goat anti-mouse IgM, conjugated with DyLight549 fluorophore. Images were taken with ZOE^TM^-fluorescent cell imager (Bio-Rad). For transmission electron microscopy animals were perfused with 0.9% w/v NaCl and subsequently with electron microscopy grade fixative buffer containing: 3% w/v glutaraldehyde, 3% w/v paraformaldehyde in phosphate buffer, pH 7.4 (Electron Microscopy Science). Tissues were stored in said buffer and processed as described previously^46^. In short, tissues were post-fixed with 2% osmium tetroxide and dehydrated through a graded ethanol series. The samples were subsequently cleared in propylene oxide, embedded in EPON 812 (Serva Electrophoresis GmbH) and finally cured for 20 h at 45 °C followed by additional 24 h at 60 °C. The resulting blocks were trimmed and sectioned into 60 nm thin slices with a Reichert-Jung Ultracut E ultramicrotome using a diamond knife (DiATOME Electron Microscopy Sciences). Silver-appearing sections were placed on a 150 µm mesh copper/rhodium grid (Plano GmbH). Samples were contrasted using alcoholic uranyl acetate and lead citrate. The sections were imaged using a Libra 120 transmission electron microscope (Carl Zeiss NTS GmbH) equipped with a SSCCD camera system (TRS, Olympus).

### 2.8 Protein digestion and purification

Protein digestion and DiGly peptide enrichment was performed as previously described^47,48^. Briefly, the gastrocnemius was lysed in Sodium deoxycholate (SDC) buffer (1% w/v SDC in 100 mM Tris-HCl, pH 8.5) and boiled for 5 min at 95 °C while shaking at 1000 rpm. Lysates were sonicated for 15 min (Bioruptor, Diagenode, cycles of 30 sec) and protein concentrations were determined using the Pierce BCA Protein Assay (Thermo). Chloroacetamide (CAA) and Tris(2-carboxyethyl)phosphine (TCEP) (final concentrations: 40 mM and 10 mM respectively) were added to 5 mg Protein. Samples were incubated for 10 min at 45 °C in the dark, shaking at 1000 rpm. Trypsin (1:50 w/w, Sigma) and Lysyl Endopeptidase^®^ (LysC) (1:50 w/w, Wako) were added and samples were digested overnight at 37 °C while shaking at 1000 rpm. For proteome analysis, sample aliquots (∼15 µg) were desalted in 3-layered styrenedivinylbenzene-reverse phase separation (SDB-RPS) StageTips (Empore). Briefly, samples were diluted with 2% Trifluoroacetic acid (TFA) in isopropanol (1:1) to a final volume of 200 µL. Thereafter, samples were loaded onto StageTips and sequentially washed with 200 µL of 1% TFA in isopropanol and 200 µL 0.2% TFA/2% Acetonitrile (ACN). Peptides were eluted with freshly prepared 60 µL of 1.25% ammonium hydroxide (NH_4_OH)-80 % ACN and dried using a SpeedVac centrifuge (Thermo). Dried peptides were resuspended in 6 µL 2% ACN/0.1% TFA. DiGly enrichment samples were diluted with 1% TFA in isopropanol (1:5). For peptide cleanup SDB-RPS cartridges (Strata™-X-C, 200 mg/6 mL, Phenomenex Inc.) were equilibrated with 8 bed volumes (BV) of 30% MeOH/1% TFA and washed with 8 BV of 0.2% TFA. Samples were loaded by gravity flow and sequentially washed twice with 8 BV 1% TFA in isopropanol and once with 8 BV 0.2% TFA-2% ACN. Peptides were eluted in two steps with 4 BV 1.25% NH_4_OH-80% ACN each and diluted with ddH_2_O to a final ACN concentration of 35% ACN. Subsequently, samples were dried via SpeedVac centrifuge overnight.

### 2.9 Diglycine (DiGly) peptide enrichment

For DiGly peptide enrichment the PTMScan® Ubiquitin Remnant Motif (K-ε-GG) (Cell Signaling) was used. The peptides were resuspended in immunoaffinity purification (IAP) buffer and sonicated (Bioruptor, Diagenode) for 15 min Protein concentration was determined by Pierce BCA Protein Assay (Thermo). Antibody coupled beads were cross-linked like previously described^48^. Briefly, one vial of antibody coupled beads was washed with cold cross-linking wash buffer (2000 *g*, 1 min). Subsequently the beads were incubated in 1 mL cross-linking buffer at room temperature for 30 min under gentle agitation. The reaction was stopped by washing twice with 1 mL cold quenching buffer (200 mM ethanolamine, pH 8.0) and incubating for 2 h in quenching buffer at room temperature under gentle agitation. Cross-linked beads were washed with cold IAP buffer and used directly for DiGly peptide enrichment. For DiGly enrichment 3 mg of peptide is used with 1/8 of a vial of cross-linked antibody beads. Peptides were added to the cross-linked beads and the volume was adjusted to 1 mL with IAP buffer and incubated for 2 h at 4 °C under gentle agitation. The beads were washed with cold IAP buffer and ddH_2_O. The enriched peptides were eluted by adding 200 µL 0.2% TFA onto the beads and shaking 5 min at 1400 rpm. After centrifugation, the supernatant was transferred to SDB-RPS StageTips and the peptides washed, eluted, and dried as previously described for total proteome samples.

### 2.10 LC-MS/MS proteome and ubiquitome measurements

Liquid chromatography of total proteome and ubiquitome samples was performed on an EASY-nLC^TM^ 1200 (Thermo) with a constant flow rate of 10 µL/min and a binary buffer system consisting of buffer A (0.1% v/v formic acid) and buffer B (80% v/v acetonitrile, 0.1% v/v formic acid) at 60 °C. The column was in-house made, 50 cm long with 75 µm inner diameter and packed with C18 ReproSil (Dr. Maisch GmbH, 1.9 µm). The elution gradient for the ubiquitome started at 5% buffer B, increasing to 25% after 73 min, 50% after 105 min and 95% after 110 min. The gradient for the total proteome started at 5% buffer B and increased to 20% after 30 min, further increased at a rate of 1% per minute to 29%, following up to 45% after 45 min and to 95% after 50 min. The MS measurement was performed on a Thermo Exploris 480 with injection of 500 ng peptide. Fragmented ions were analyzed in Data Independent Acquisition (DIA) mode with 66 isolation windows of variable sizes. The scan range was 300 – 1650 *m/z* with a total scan time of 120 min, an Orbitrap resolution of 120,000 and a maximum injection time of 54 ms. MS2 scans were performed with a higher-energy collisional dissociation (HCD) of 30 % at a resolution of 15,000 and a maximum injection time of 22 ms. The MS measurement of the total proteome was performed equivalently, yet including High-Field Asymmetric Waveform Ion Mobility (FAIMS) with a correction voltage of −50 V and a total scan time of 60 min. The injection time for the full scan was 45 s and the MS2 injection time was set to 22 s.

### 2.11 Proteome and Ubiquitome data analysis

DIA raw files were processed using Spectronaut (13.12.200217.43655) in directDIA mode. The FASTA files used for the peptide identification were: Uniprot Mus Musculus (10.03.2022) with 21989 entries, Uniprot Mus Musculus isoforms (10.03.2022) with 41771 entries and MaxQuant Contaminants for filtering (245 entries). Protein groups in the total proteomes were quantified through label free quantification, following the MaxLFQ algorithm. Analysis was performed via Perseus (version 1.6.2.3). The site tables for ubiquitome samples were created with the Perseus plugin “Peptide Collapse” (version 1.4.2)^49^. Values were log2-transformed and missing values were replaced by imputation from normal distribution with a width of 0.3 and downshift 1.8 separately for each sample. Ubiquitome was normalized to total proteome. Comparison between conditions of proteome and ubiquitome was performed via Student’s *t*-test in R version 4.2.2 (*P*-value cutoff 0.1 and 0.2 for GO-analysis).

### 2.12 Non-targeted metabolomics, data processing, and analysis

Untargeted metabolomics of muscle and plasma was performed by the Metabolomics and Proteomics Core facility (MPC) at Helmholtz Munich (Neuherberg, Germany) as previously described.^50^ Briefly, frozen muscle tissue was homogenized at 4 °C with 1.4 mm ceramic beads in water (5 µL/mg). To precipitate the protein and extract metabolites, 500 µL MeOH extraction solvent containing recovery standard compounds was added to each 100 µL of plasma or tissue homogenate. Supernatants were aliquoted and dried under nitrogen stream (TurboVap 96, Zymark) and stored at –80 °C until measurement. An ultra-high performance liquid chromatography-tandem mass spectrometry (UPLC-MS/MS) based analytical platform licensed by Metabolon (Durham, NC, USA) using Acquity UPLC (Waters) coupled to a Q Exactive mass spectrometer (Thermo) was used to measure the samples. Three separate UPLC-MS/MS injections were performed: two injections for analysis by UPLC-MS/MS positive ionizations, e.g., early, and late eluting compounds, and one injection for analysis by UPLC-MS/MS negative ionization. The resulting MS/MS data were matched to Metabolon’s proprietary chemical library, which includes retention time, molecular weight (m/z), preferred adducts, and in-source fragments, as well as associated MS/MS spectra for all molecules in the library. Peak area counts were median-normalized. Metabolites with more than 50 % missing values were excluded, and missing values were imputed by k-nearest-neighbor algorithm. Values were log10 transformed for univariate and statistical analyses.

### 2.13 Seahorse

Seahorse analysis was performed as described previously^51^. Briefly, gastrocnemii were removed and homogenized in 1 mL MIBI buffer (210 mM d-Mannitol (Sigma), 70 mM sucrose (Sigma), 5 mM HEPES (Roth), 1 mM EGTA (Roth), 0.5% w/v fatty acid-free BSA (Sigma), pH 7.2) via douncer (Carl Roth). The homogenate was centrifuged at 800 *g* for 10 min at 4 °C. The supernatant was centrifuged at 8000 *g* and the pellet washed twice more with MIBI buffer. The pellet was resuspended in MASI buffer (220 mM d-Mannitol (Sigma), 70 mM sucrose (Sigma), 10 mM KH_2_PO_4_ (Sigma), 5 mM MgCl_2_ (Sigma), 2 mM HEPES (Roth), 1 mM EGTA (Roth), 0.2% w/v fatty acid-free BSA (Sigma), pH 7.2 at 37 °C) with 10 mM succinate (Sigma) and 2 mM pyruvate (Agilent) and the mitochondria concentration assessed by Pierce BCA Protein Assay (Thermo). The assay was performed on a Seahorse XF24-analyzer (Agilent) with 1 µg mitochondria, 16 mM ADP (Sigma), 1 µM Oligomycin (Sigma), 40 µM FCCP (Sigma), 25 µM Rotenone/AntimycinA (Sigma). Injections and mix/measurement cycle times for the assay are illustrated in Supplementary Table 3.

### 2.14 Oroboros

Mitochondrial respiration was measured in permeabilized skeletal muscle fibers using a high resolution respirometer (Oxygraph-2k, Oroboros Instruments) as previously described.^52^ Briefly, all respiratory measurements were made in duplicates in a hyperoxygenated environment ([O_2_], 450-200 nmol/mL) to avoid potential oxygen limitations of the respiration. Skeletal muscle fibers were permeabilized with saponin (50 µg/mL) for 30 min in BIOPS buffer followed by 2 × 10 min wash in respiration medium 5 (MiR05). Leak respiration was assessed after addition of 2 mM malate, 10 mM glutamate and 5 mM pyruvate. Next, NADH-linked respiration was measured by adding 5 mM ADP and 3 mM MgCl_2_ followed by 10 mM Cytochrome C to evaluate outer membrane integrity. Oxidative phosphorylation capacity with simultaneous electron input of complex I and II was assessed with 10 mM succinate. Lastly, maximum respiration rate was determined by adding 1 mM FCCP. All respiratory rates were measured in MiR05 buffer (containing 110 mmol/L sucrose, 60 mmol/L potassium lactobionate, 20 mmol/L Hepes, 10 mmol/L KH_2_PO_4_, 1 g/L BSA, 0.5 mmol/L EGTA, 20 mmol/L taurine, 3 mmol/L MgCl_2_·6H_2_O, pH 7.1) at 37 °C.

### 2.15 Flow cytometry analysis

Circulating pro-and anti-inflammatory cytokines and chemokines in plasma were analyzed using the LEGENDplex™ Mouse Anti-Virus Response Panel (Biolegend). Analysis was performed with MACSQuant® Analyzer 10 (Miltenyi). To determine immune cell infiltration in muscle tissue the gastrocnemius was harvested and minced and added to a digestion buffer with 2 mg/mL collagenase type 2 and 50 µg/mL DNaseI in DMEM Glutamax medium (Thermo) with 10% FBS (Sigma) and 1% Pen/Strep (Sigma). The muscle was incubated 45 min at 37 °C while shaking. After digestion the suspension was passed through a 70 µm cell strainer and centrifuged at 300 *g* for 5 min at 4 °C. The pellet was resuspended in FACS buffer (DPBS, 1% FBS, 2 mM EDTA, 0.1% sodium azide w/v). The cells were washed with cold PBS and fixed at room temperature with 4% PFA for 15 min, washed twice with PBS and resuspended in FACS buffer. For blocking of the cells FACS buffer was added with supplementation of 1:100 anti-mouse CD16/32 (Fc block, BioLegend) and cells were incubated on ice for 10 min. After blocking cells were centrifuged at 500 *g* for 5 min at 4 °C and the supernatant discarded. Cells were stained for 30 min in the dark at 4 °C with anti-CD45.2, anti-CD11b and anti-Ly6c (Supplementary Table 2). Compensation was performed using countbright beads (Invitrogen). The samples were analyzed via MACSQuant® Analyzer 10 (Miltenyi). The data was analyzed using FlowJo (v.10). Single stained samples were used to correct for spectral overlaps and Fluorescence Minus One ensured correct gating controls. The gating strategies can be found in Figure EV2. We calculated absolute cell numbers according to countbright beads (Invitrogen) instructions and normalized cell number per gastrocnemius tissue.

### 2.16 Tissue-wide expression profiling and data integration

Expression values across 43 human tissues were log-transformed to evaluate tissue-specific enrichment. This cross-tissue analysis was further contextualized using curated datasets from the Cardiovascular Disease Knowledge Portal^53^.To define the systemic transcriptional landscape of *NFE2L1*, bulk RNA-sequencing metadata were integrated from the Human Protein Atlas (HPA) and the GTEx (v8) consortium. We specifically analyzed bulk RNA-seq profiles from 13,160 myonuclei in skeletal muscle (*n* = 818 human samples) and 33,023 cardiomyocytes in heart muscle (*n* = 452 human samples). Mean expression levels were quantified using normalized counts per million (nCPM) to allow for robust inter-tissue comparison.

### 2.17 Single-fiber transcriptomics and pseudobulk re-analysis

Metabolic stratification of *NFE2L1* and *NFE2L2* was conducted via re-analysis of single-fiber RNA-seq data obtained from the ArrayExpress database (accession number E-MTAB-12529). Individual myofibers were classified into fast-twitch, intermediate, and slow-twitch clusters based on their molecular profiles. Transcriptional abundance was quantified using a pseudobulk aggregation strategy (Seurat -v4), wherein raw counts from fibers exhibiting identical physiological profiles were pooled. This approach enhanced statistical power and permitted direct comparative analysis of Nrf-family member enrichment across the metabolic spectrum of skeletal muscle fibers.

### 2.18 Re-analysis of single-nucleus RNA sequencing (snRNA-seq)

To resolve the myonuclear and interstitial heterogeneity of skeletal muscle at single-cell resolution, we performed a re-analysis of snRNA-seq data from the GEO database (accession number GSE240061). Data processing was executed using the Seurat (v4) bioinformatic pipeline. The distribution and expression density of *NFE2L1* across distinct cellular clusters were evaluated via violin plot (VlnPlot) visualizations to identify cell-type-specific transcriptional signatures.

### 2.19 GeneBridge systems genetics and functional enrichment

Functional gene-to-phenotype associations for *NFE2L1* were determined using the GeneBridge framework^54^. We utilized the Gene-Module Association Determination (G-MAD) approach, which facilitates the annotation of gene function by integrating multispecies expression compendia. This method was employed to identify Gene Ontology (GO) terms and biological pathways significantly correlated with *NFE2L1* expression specifically within skeletal muscle modules. Associations exceeding the established significance threshold of > 0.268 were considered robust. Significant functional terms were visualized using a distribution plot of GO pathways to highlight the biological axes coupled with *NFE2L1* activity.

### 2.20 Statistics

Data were analyzed with ImageLab, Microsoft Excel, GraphPad Prism and R (4.2.2). Unless otherwise specified, data are shown as mean ± standard error of the mean (SEM), including individual measurements. Multiple Student’s *t*-test was used for experiments when comparing two groups and one variable and two-way ANOVA with Bonferroni post-hoc test was used for comparing two groups with two variables. If not declared otherwise, *P*-values lower than 0.05 were considered significant and are as such indicated in the graphs with an asterisk between groups, non-significant groups have no indication. For proteomic analyses p-value cutoff was 0.2 in GO analysis and 0.1 for further analyses. For the differential expression analysis of RNAseq, we used R package *limma*^55^ to compute the *P*-values and adjust for hypothesis testing using FDR.

## Supporting information

Supplemental material (uncropped blots)

## Acknowledgement

We thank Silvia Weidner, Thomas Pitsch, Daniela Hass, and Daniel Brandt for excellent technical assistance. We are grateful to Henrika Jodeleit for their help with animal experiments. We express gratitude to Joel Guerra for his input and encouragement during mitochondria isolation. We thank Anna Artati for assistance with metabolomics and Yvonne Jansen for supporting the histology experiments. Special thanks to Gökhan Hotamışlıgil for his continuous support during the project and to Sander Kersten, Patrick Schrauwen, and Wouter van Marken Lichtenbelt for their kind encouragement and guidance. We thank all Bartelt lab members for discussions and the enjoyable atmosphere. M.R. is funded by the European Research Council (ERC) under the European Union’s Horizon 2020 research and innovation program (#949017), and Helmholtz Association - Initiative and Networking Fund. P.N. and M.P.M. thank the German Center for Diabetes Research (DZD) for funding. N.K. and D.T.H. are supported by Emmy-Noether DFG (KR5166/1-1). T.M. and Z.G.-H. were supported by funding from the Novo Nordisk Foundation Center for Basic Metabolic Research. S.H. was supported by the Deutsches Zentrum für Herz-Kreislauf-Forschung DZHK. A.B. was supported by DFG Priority Programs 2306 and 2453, the DFG CRC1123 on Atherosclerosis, DZHK/DZG, and the ERC Starting Grant PROTEOFIT. We apologize to colleagues whose work we could not cite due to space limitations.

## Data availability statement

Data sets will be uploaded upon acceptance of the article.

## Author contributions

I.L. designed and performed experiments, analyzed data, and wrote the manuscript. C.J., L.M., N.W., A.M.G., E.G., I.T., and M.R. performed experiments and analyzed data. F.R., S.Kh., S.Ko. and S.H. analyzed data. D.H. and N.K. performed and analyzed proteomics data. D.T.E. and J.W. performed electron microscopy. S.L., T.M. and Z.G.-H. performed Oroboros measurements and analysis. K.D. performed and analyzed metabolomics data. A.B. made coffee, designed experiments, analyzed data, and wrote the manuscript. All authors read and commented on the manuscript.

---

**Supplementary Table 1:**
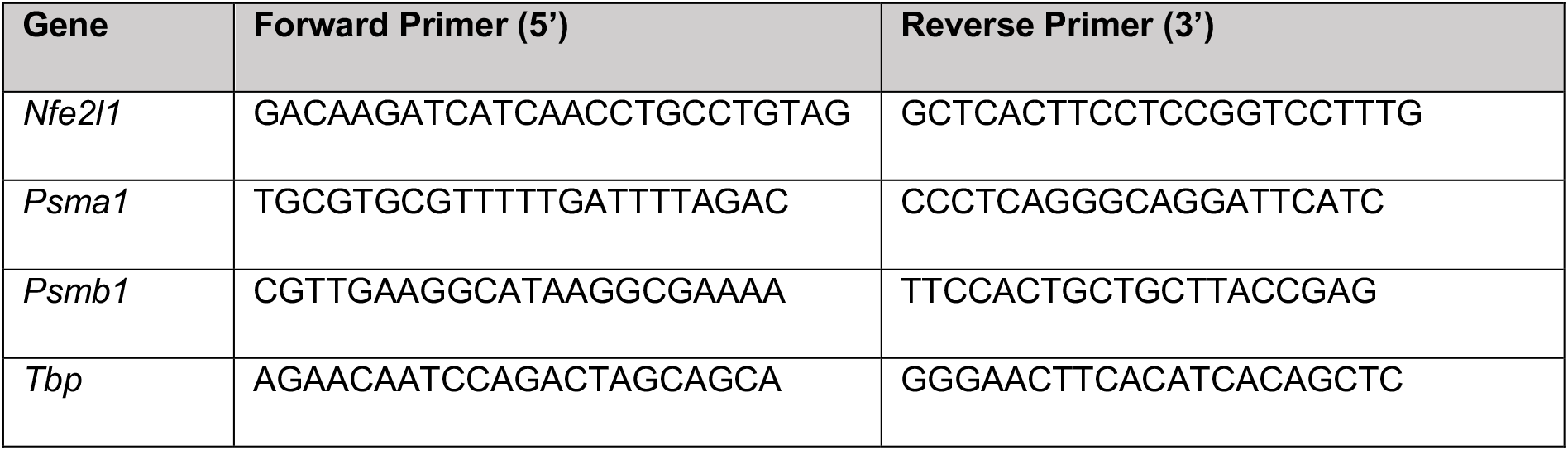

**Supplementary Table 2:**

| Antibody | Company (Cat. No.) | Dilution |
| --- | --- | --- |
| Gapdh | Cell signaling technologies (5174) | 1:1000 |
| LC3B | Cell signaling technologies (2775) | 1:1000 |
| Nfe2l1 | Cell signaling technologies (D5B10) | 1:500 |
| Total OXPHOS rodent | Abcam (ab110413) | 1:1000 |
| p62 | Abcam (ab155686) | 1:1000 |
| Ubiquitin (P4D1) | Cell signaling technologies (3936) | 1:1000 |
| β-tubulin | Cell signaling technologies (2146) | 1:1000 |
| Psmb5 | Cell signaling technologies (12919) | 1:1000 |
| Psmb6 | Cell signaling technologies (13267) | 1:1000 |
| Psmb7 | Cell signaling technologies(13207) | 1:1000 |
| Psme1 (PA28 alpha) | Cell signaling technologies (2408) | 1:1000 |
| Anti-rabbit IgG, HRP-linked | Cell signaling technologies (7074) | 1:10000 |
| Anti-mouse IgG, HRP-linked | Cell signaling technologies (7076) | 1:10000 |
| Purified anti-mouse CD16/32 Antibody (Fc Block) | BioLegend (101302) | 1:100 |
| Cd45.2 Monoclonal Antibody PerCP-Cyanine5.5 | eBioscience (45-0454-82) | 1:100 |
| Cd11b Monoclonal Antibody APC-eFluor 780 | eBioscience (45-0112-82) | 1:100 |
| Ly-6C Monoclonal Antibody Brilliant Violet 510 | BioLegend (128033) | 1:50 |

**Supplementary Table 3:**

| Command | Time |
| --- | --- |
| Baseline | Wait: 00:10:00<br>Mix: 00:01:00<br>Wait: 00:03:00<br>Duration: 00:14:00 |
| Measurement Baseline | Mix: 00:01:00<br>Wait: 00:00:00<br>Measure: 00:03:00<br>Cycles: 2<br>Duration: 00:08:00 |
| ADP (Port A) | Mix: 00:01:00<br>Wait: 00:00:00<br>Measure: 00:04:00<br>Cycles: 1<br>Duration: 00:05:00 |
| Oligo (Port B) | Mix: 00:01:00<br>Wait: 00:00:00<br>Measure: 00:03:00<br>Cycles: 1<br>Duration: 00:04:00 |
| FCCP (Port C) | Mix: 00:01:00<br>Wait: 00:00:00<br>Measure: 00:03:00<br>Cycles: 1<br>Duration: 00:04:00 |
| Rot/AA (Port D) | Mix: 00:01:00<br>Wait: 00:00:00<br>Measure: 00:03:00<br>Cycles: 2<br>Duration: 00:08:00 |

**Supplementary Table 4:**
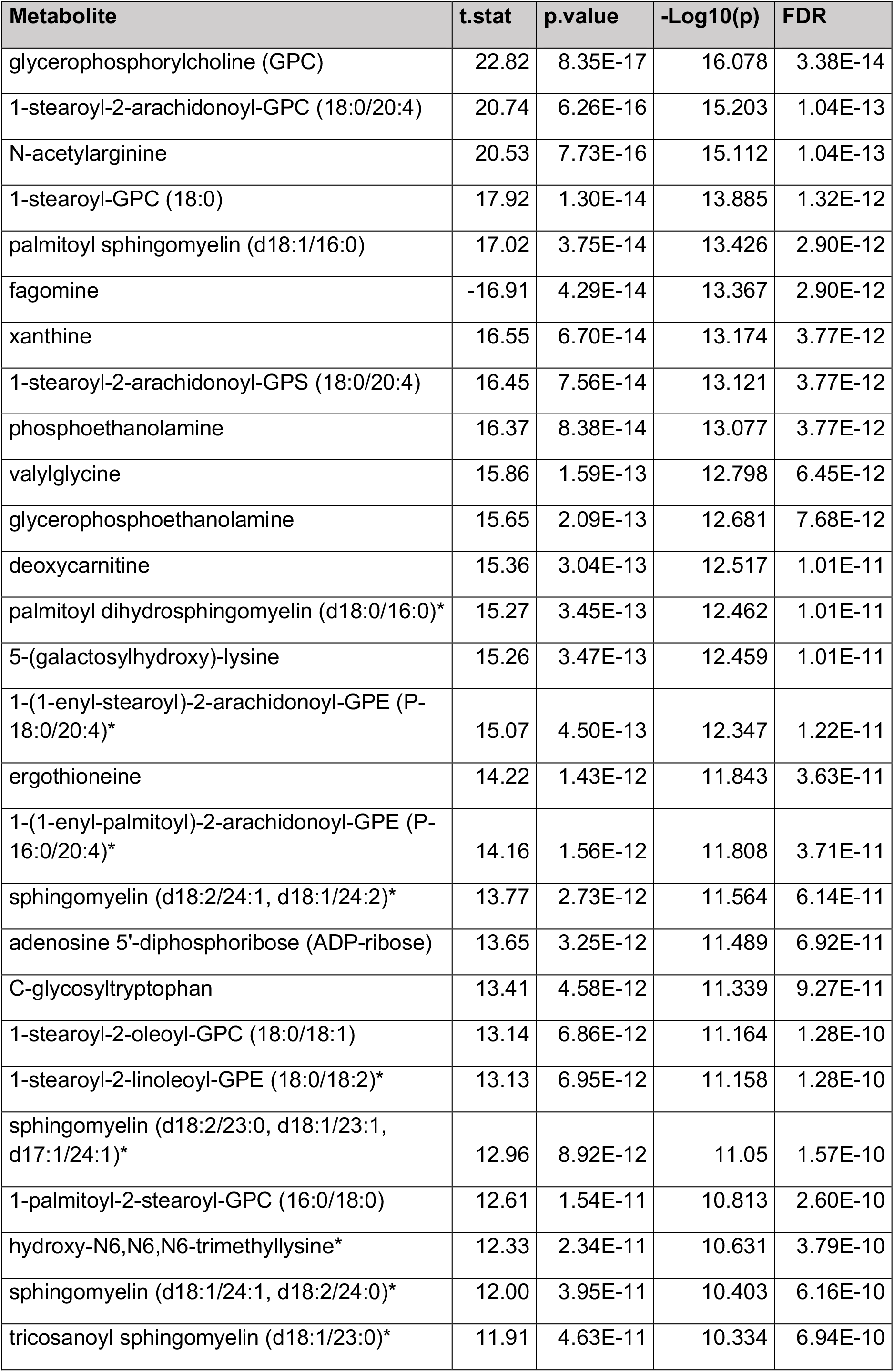
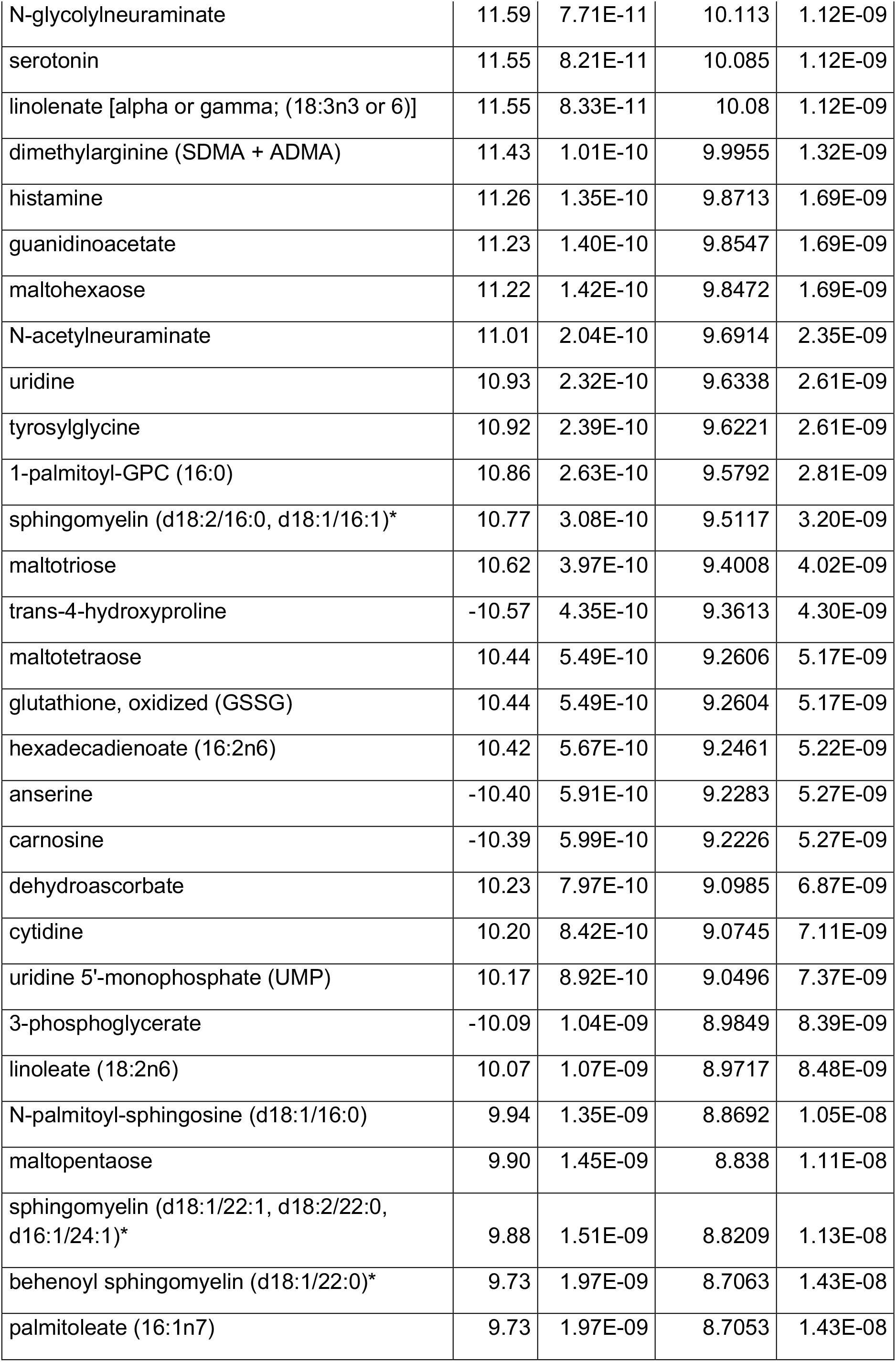
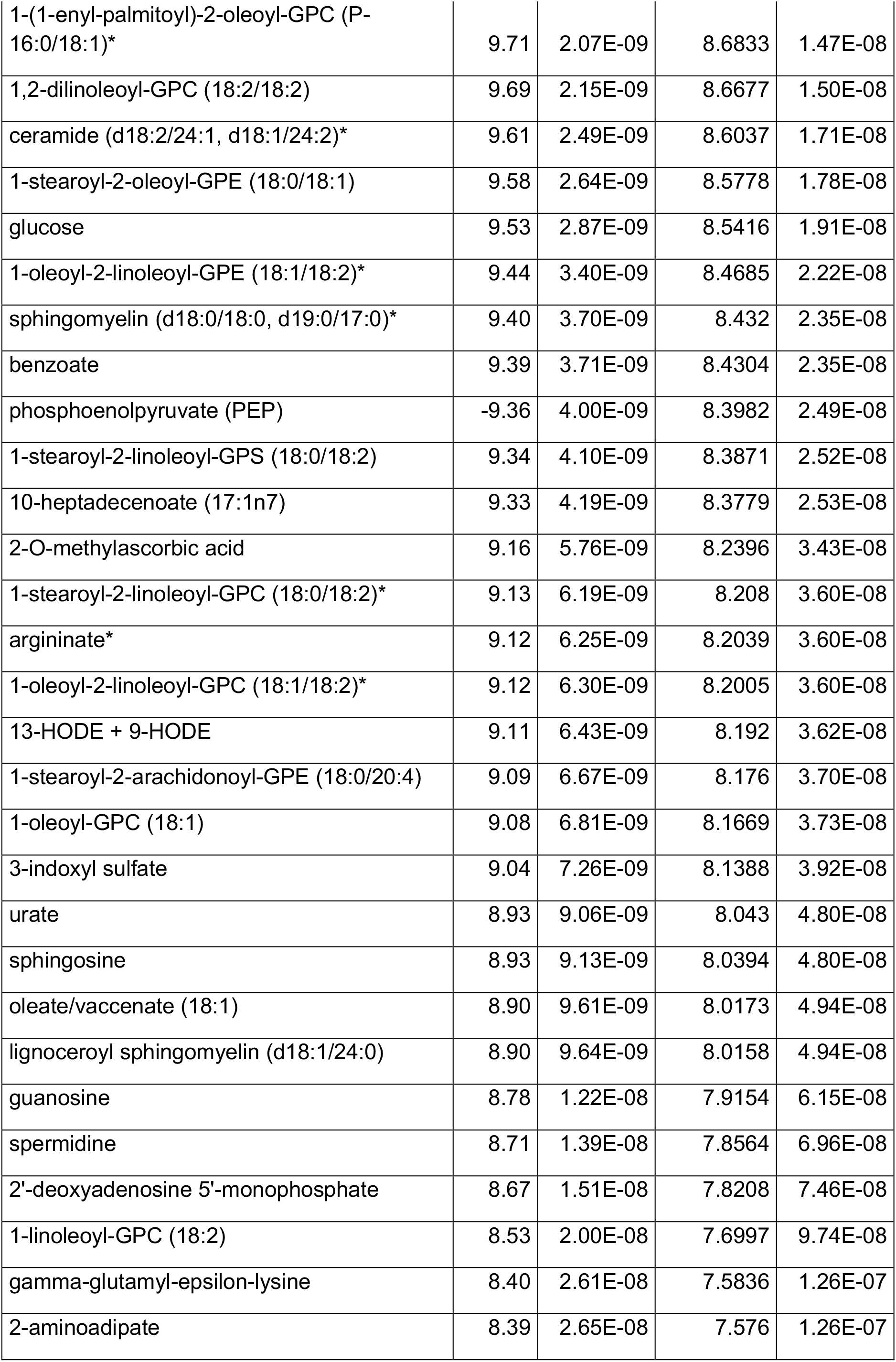
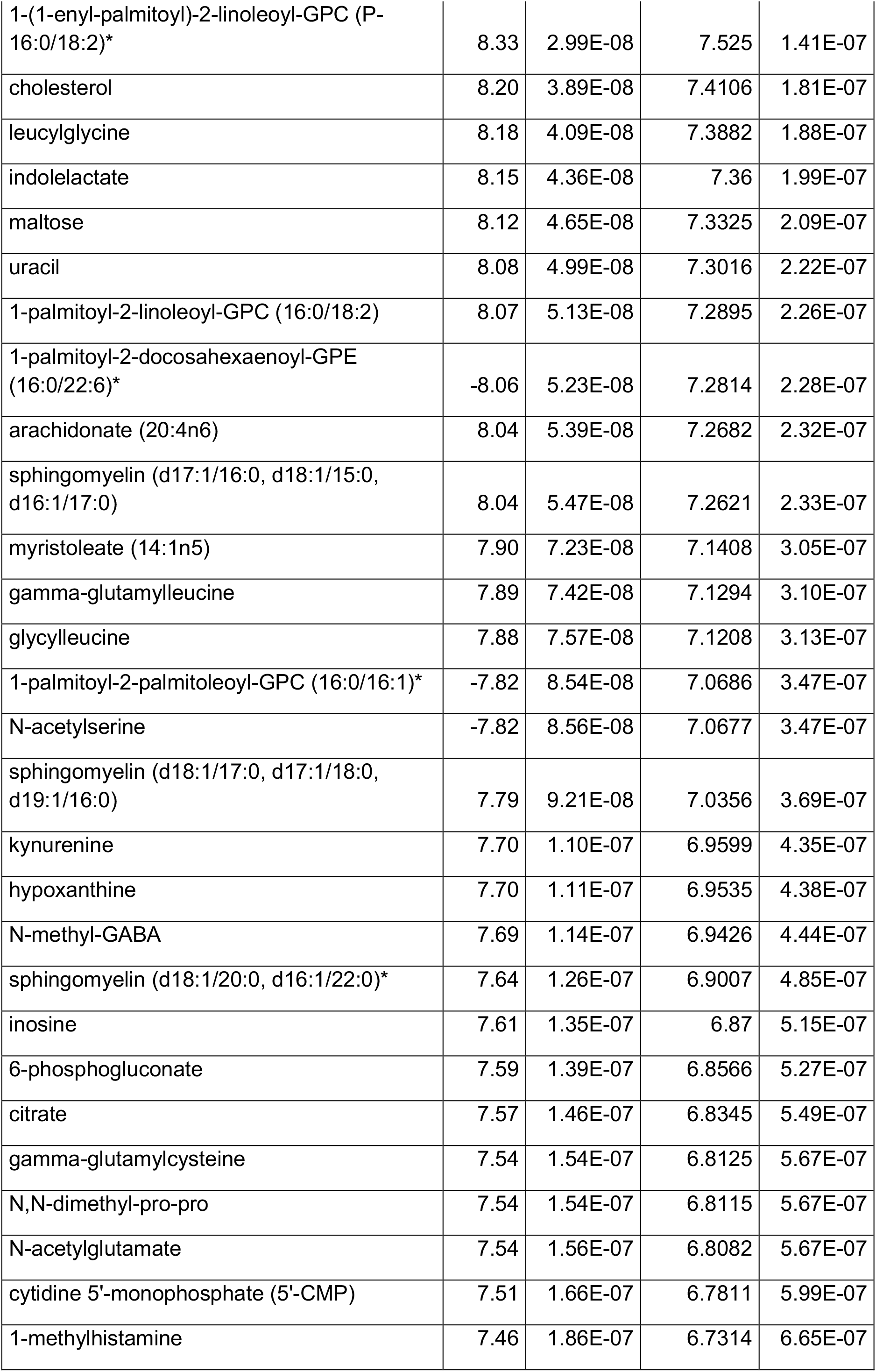
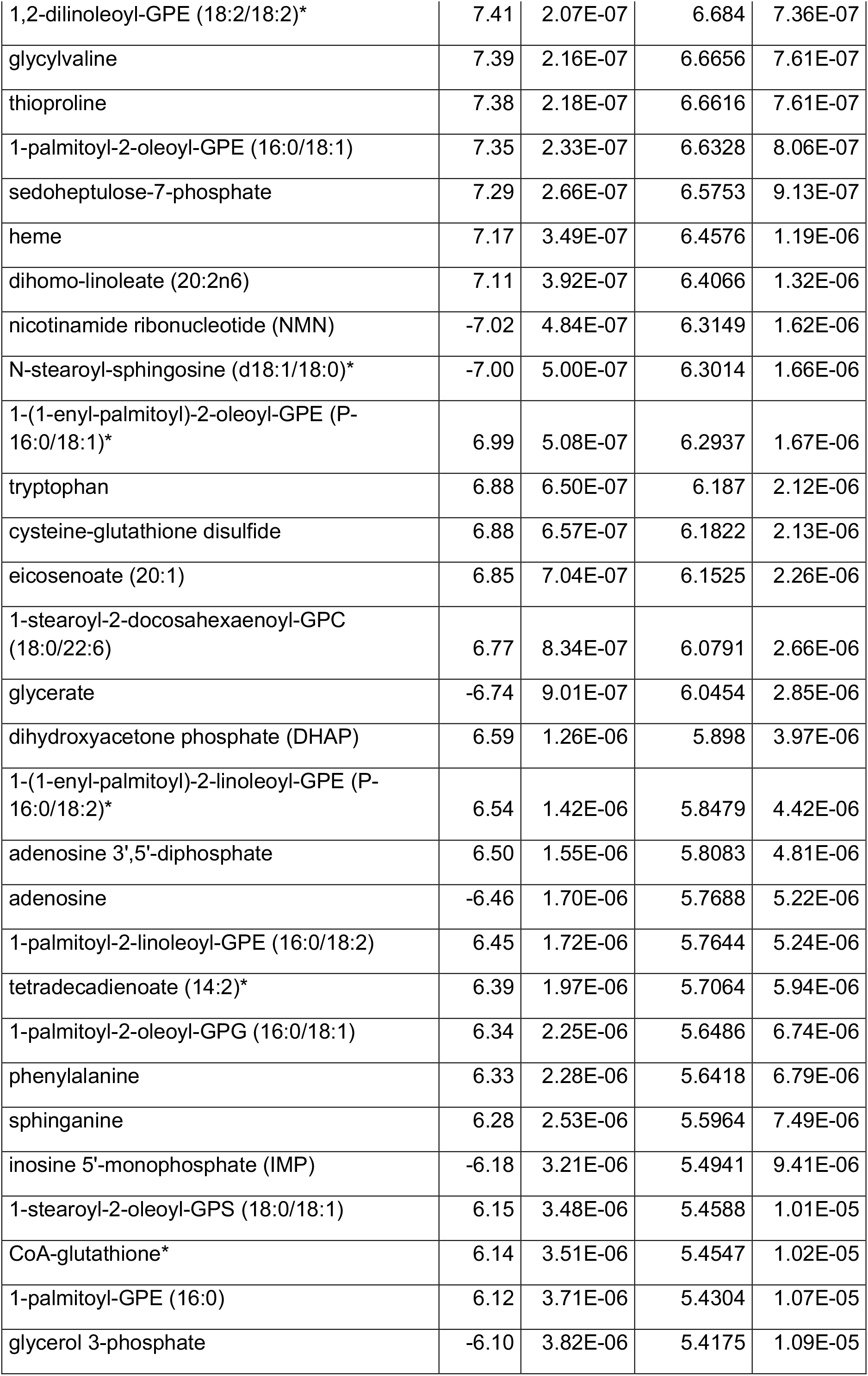
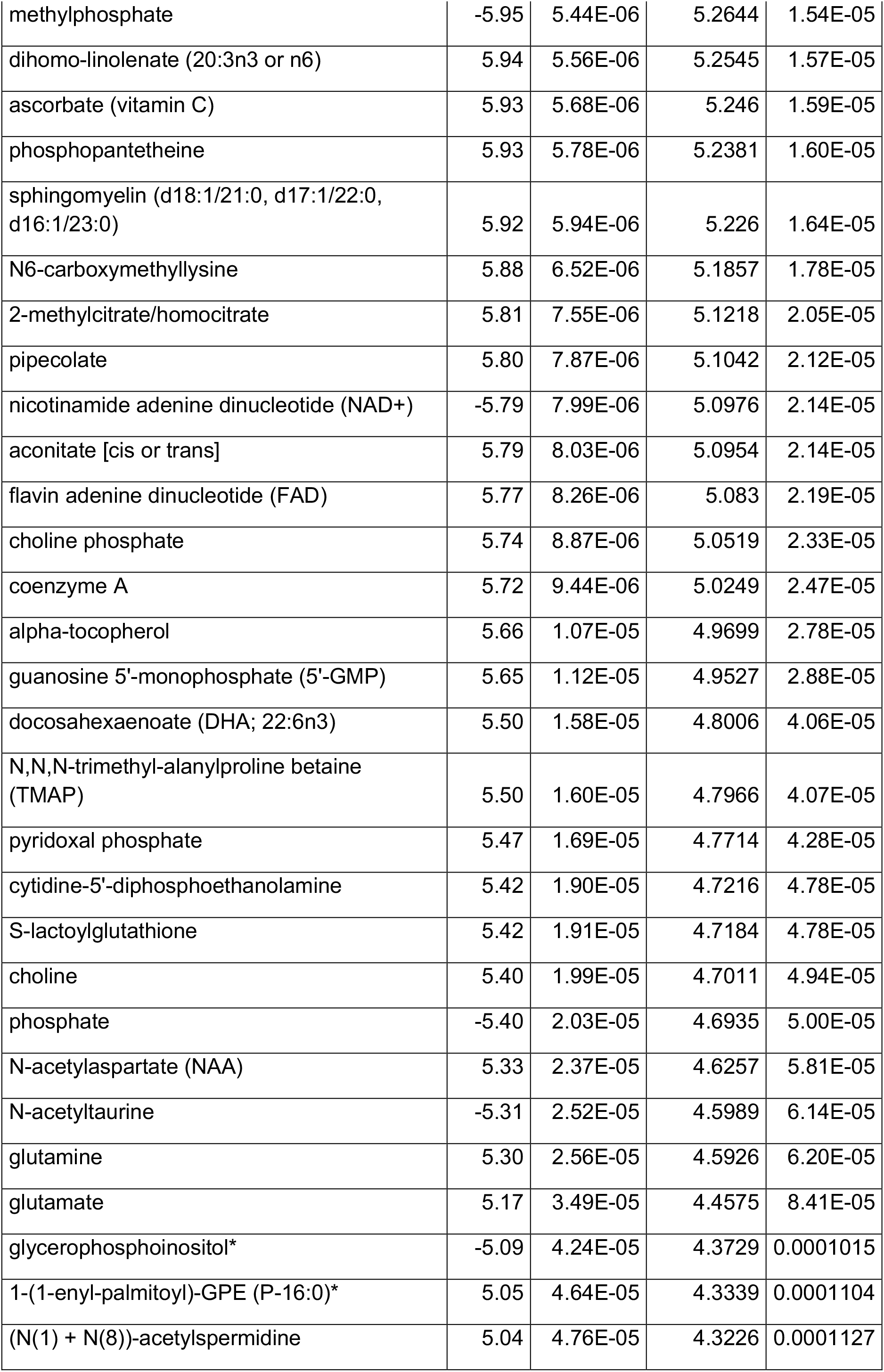
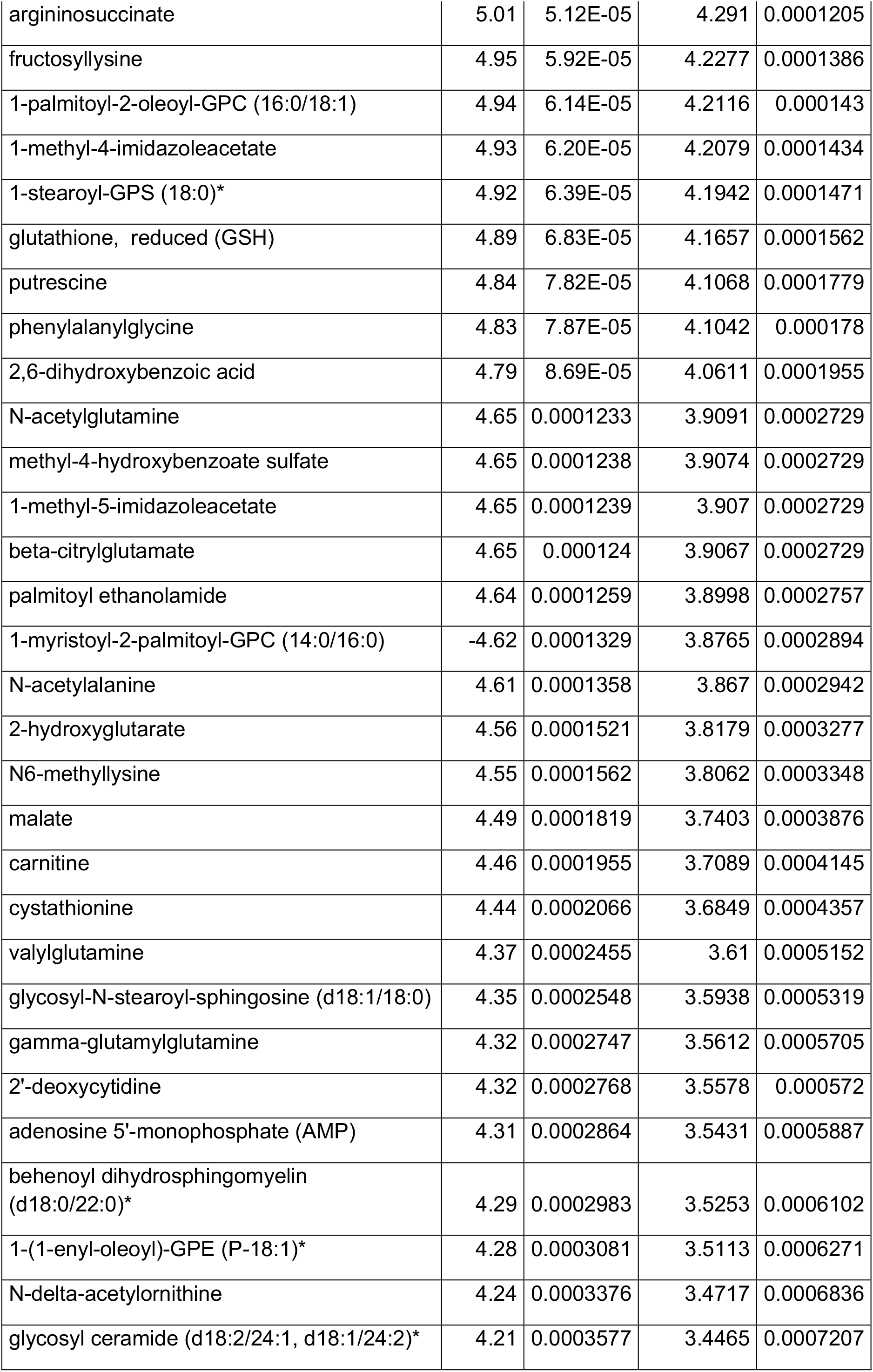
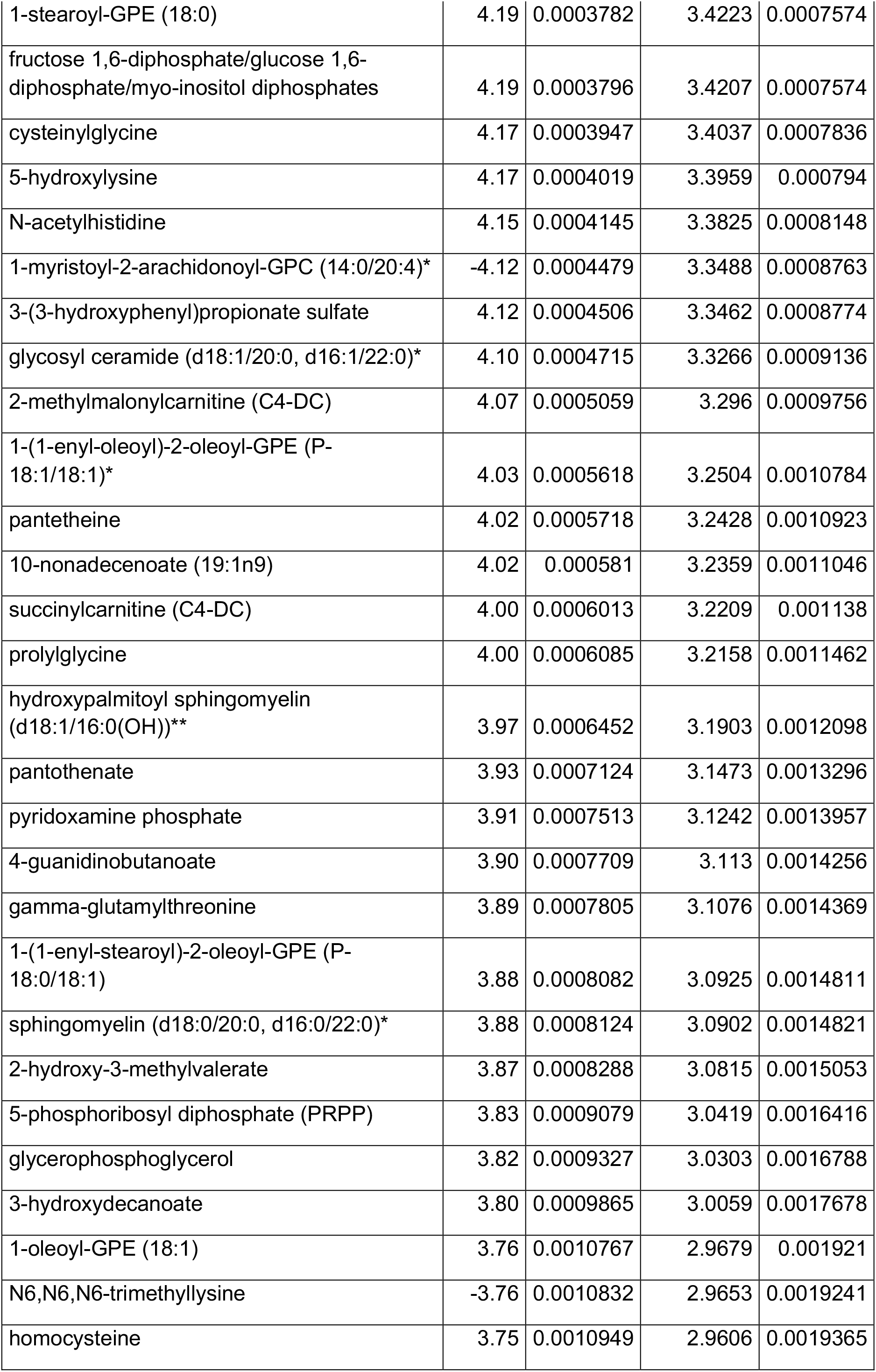
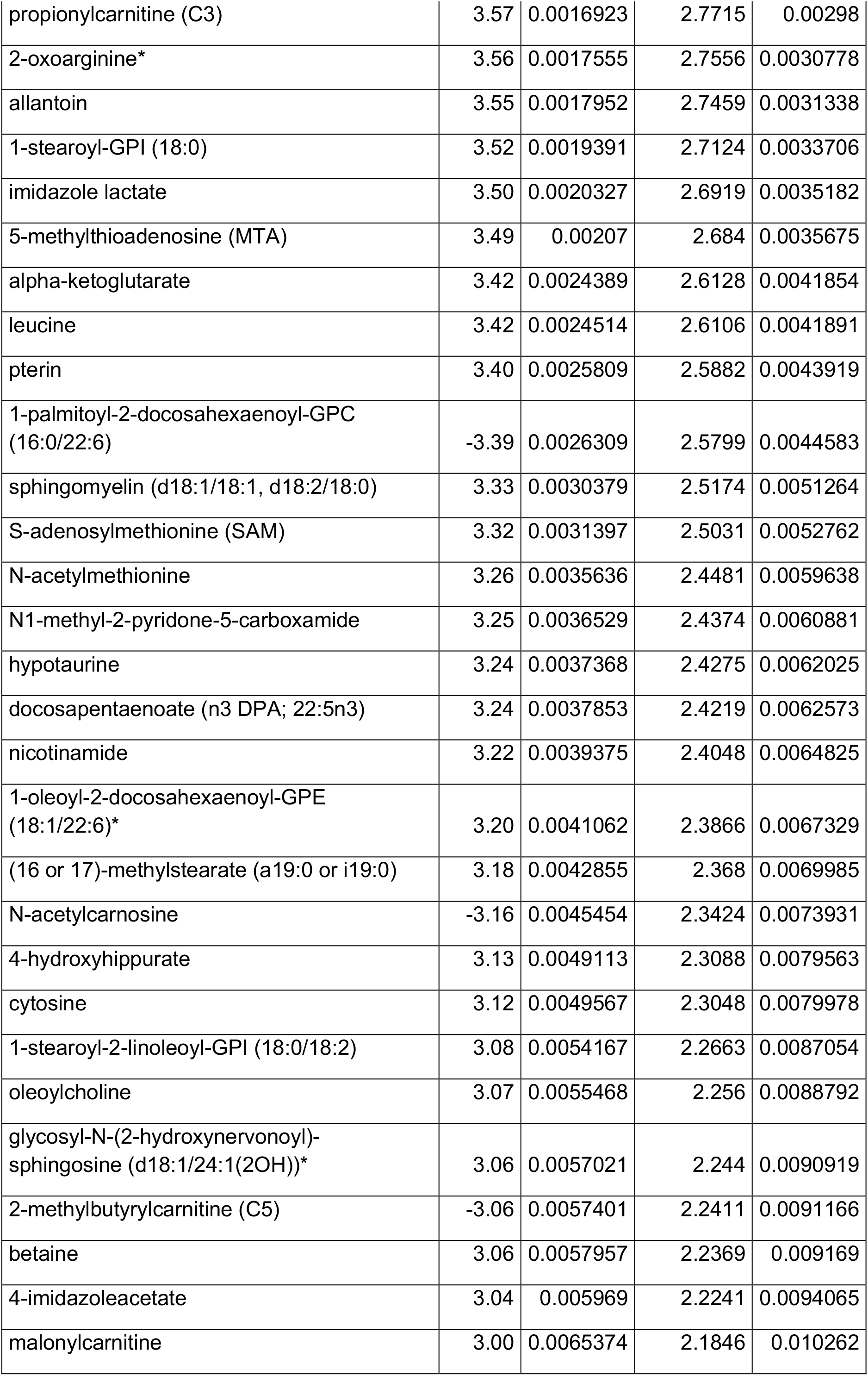
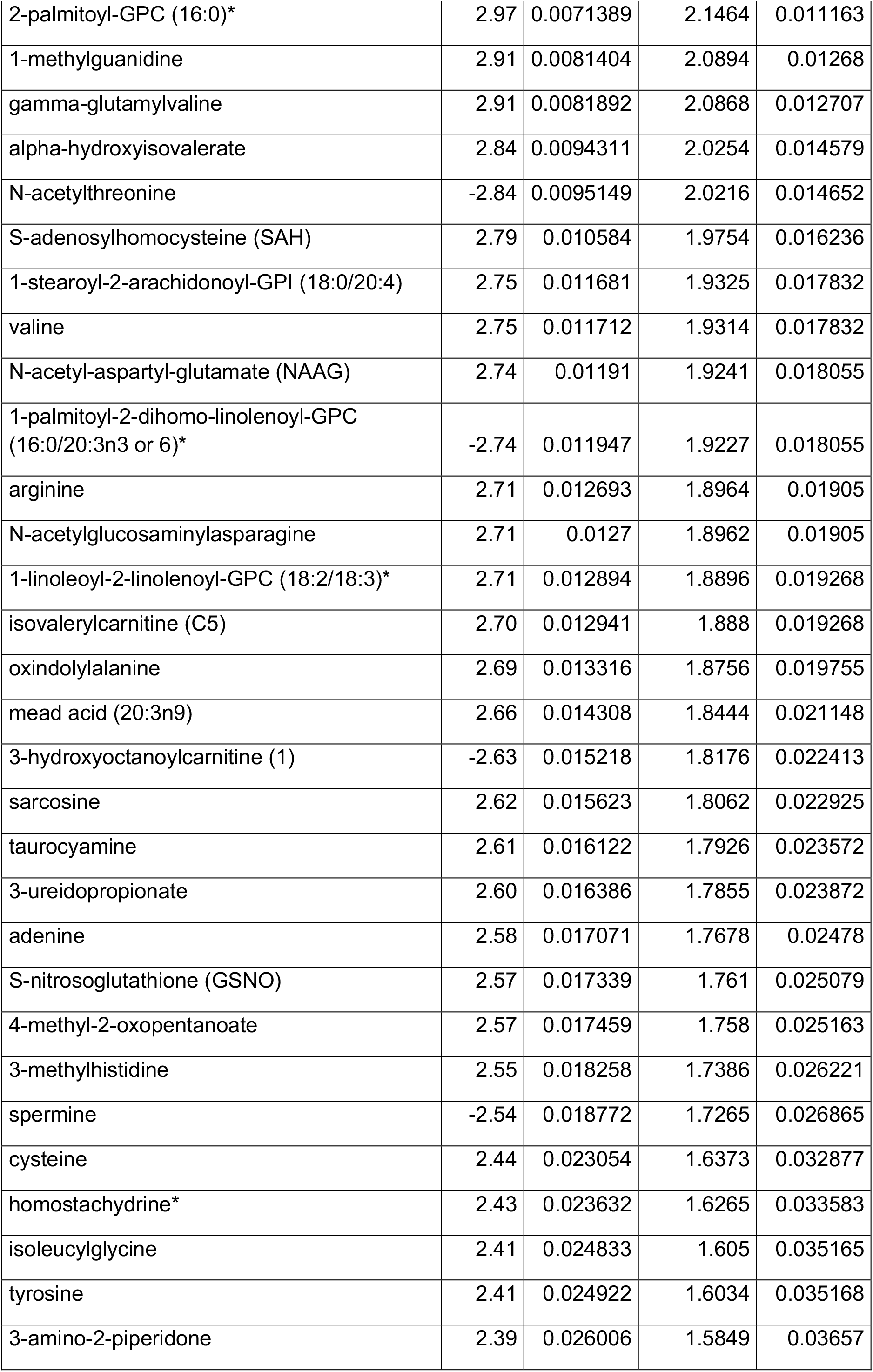
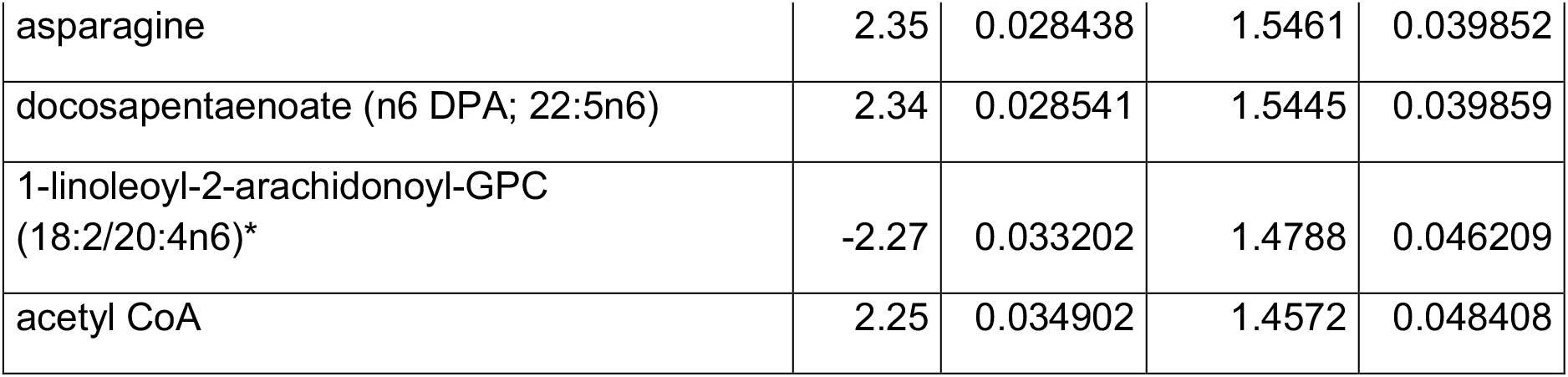

