## Supplemental figures for "The myocyte Nfe2l1-ubiquitin-proteasome system controls muscle fiber type and obesity-induced insulin resistance"

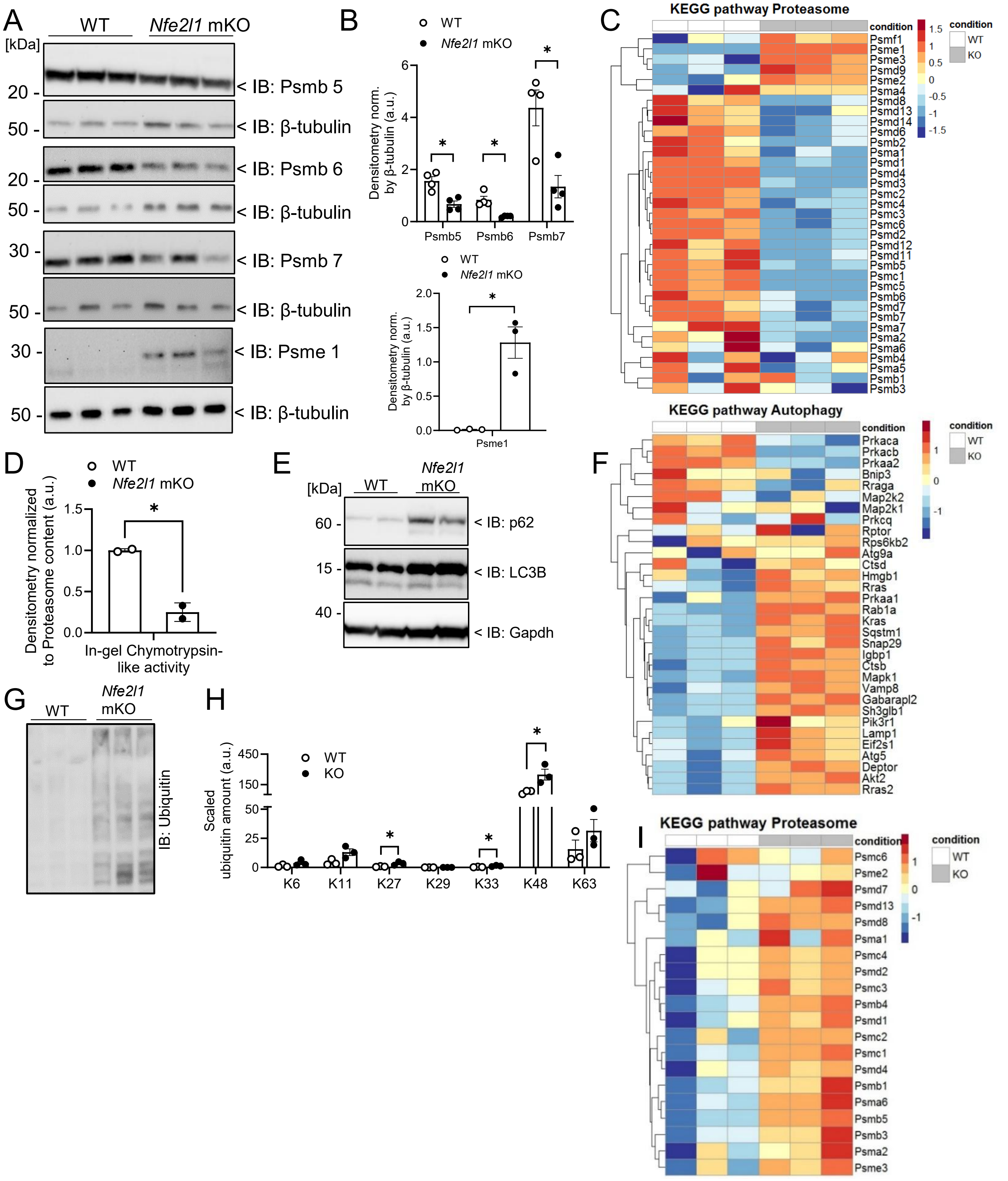


**Figure EV1: Loss of Nfe2l1 impacts proteasome and ubiquitylation landscape as well as autophagy.** (A-I) Assessment of GC tissue from *Nfe2l1* mKO mice and WT controls (*n* = 2 - 4 mice per group) via immunoblot or proteomics and ubiquitomics analyzed via mass spectrometry. (A) Immunoblot and corresponding (B) Densitometry of proteasome subunits. (C) KEGG analysis (03050) of proteasome from total proteome. (D) Densitometry corresponding to Nu-PAGE proteasome activity. (E) Immunoblot of autophagy markers. (F) KEGG analysis (4140) of autophagy from total proteome. (G) Immunoblot of ubiquitylated proteins. (H) Relative amount of ubiquitin lysine (K)-linkages. (I) KEGG analysis (03050) of proteasome from ubiquitome. The data are mean ±SEM, P<0.05 by *t*-test (B bottom, D) or 2-way ANOVA (B top, H).

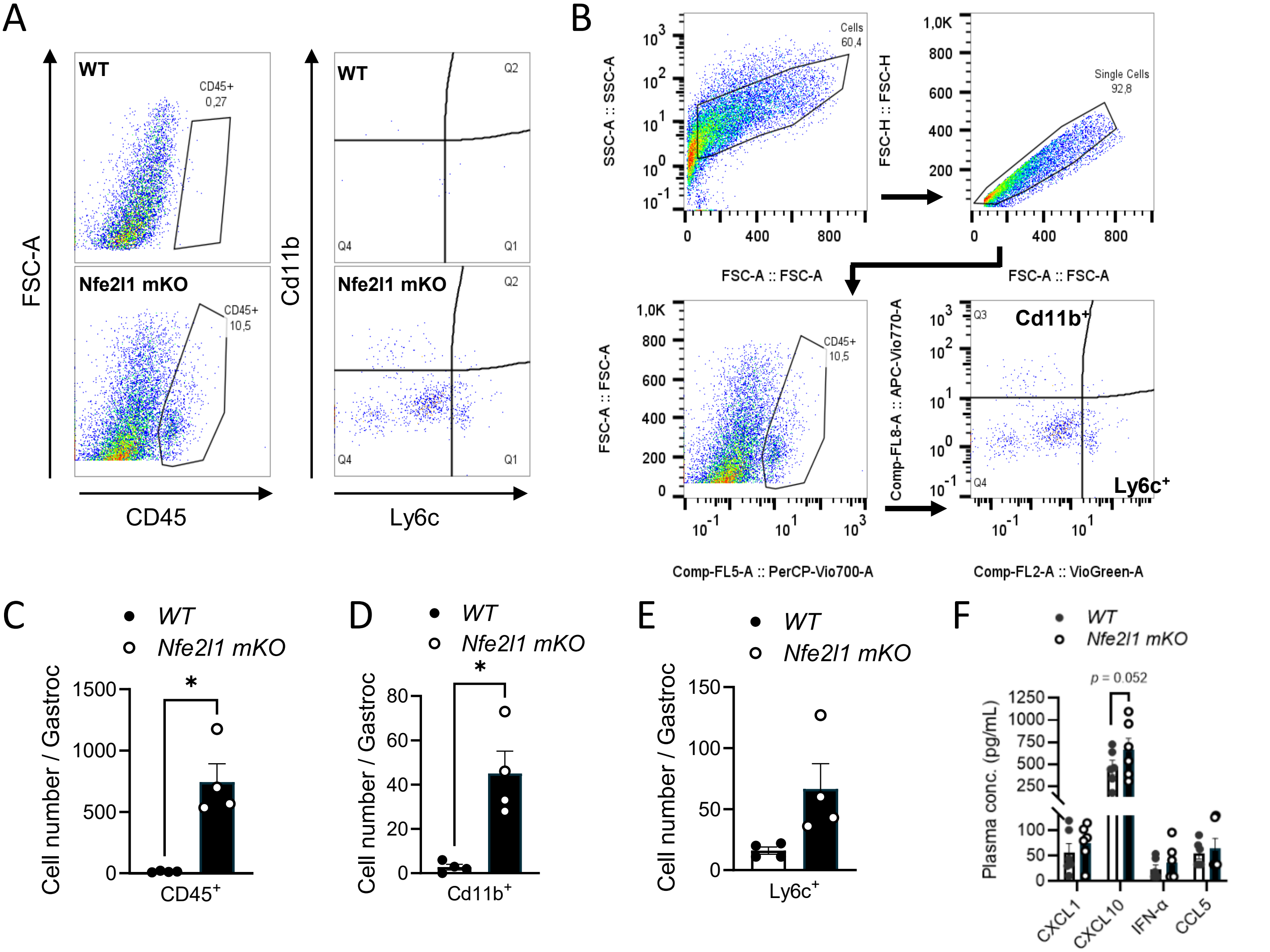


**Figure EV2: Loss of Nfe2l1 leads to immune cell infiltration in gastrocnemius muscle.**
(A) Representative flow cytometry plot of gastrocnemius from WT and *Nfe2l1* mKO mice stained for CD45, Cd11b and Ly6c. (B) Gating strategy for $CD45^+$ cells and for $Cd11b^+$ as well as $Ly6c^+$ cells within $CD45^+$ population. (C-E) Absolute cell numbers of CD45, Cd11b and Ly6c positive cells respectively ($n$ = 4 mice per group) (F) Circulating cytokines measured in plasma from WT and *Nfe2l1* mKO animals via flow cytometry ($n$ = 6 mice per group). The data are mean ±SEM, P<0.05 by *t*-test (B-D) or 2-way ANOVA (E).

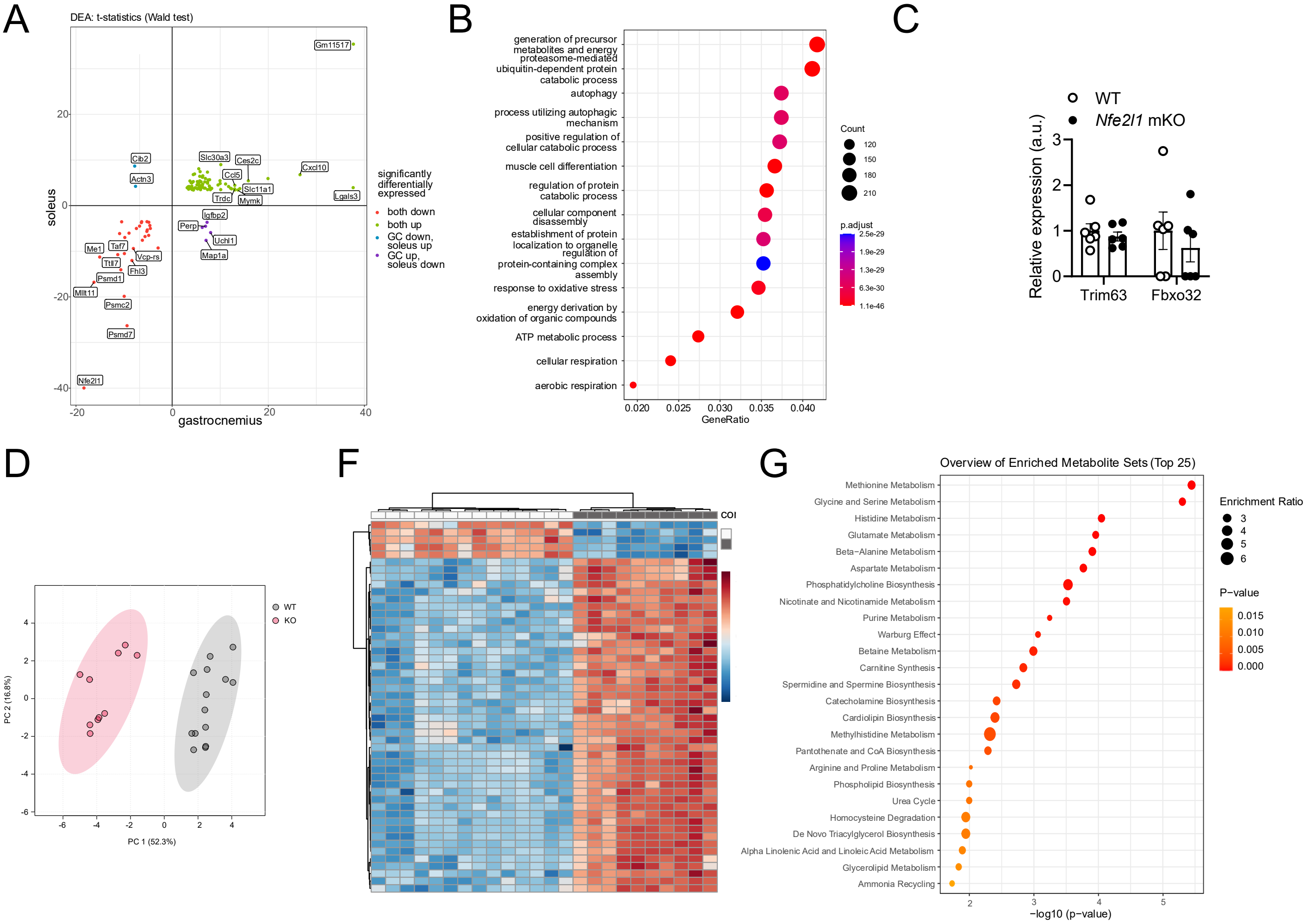


**Figure EV3: Loss of Nfe2l1 impacts catabolic processes on transcriptional level and whole-body metabolic landscape.** (A-B) RNA-seq of gastrocnemius (GC) and soleus muscle (*n* = 6 mice per group) from WT and *Nfe2l1* mKO mice. (A) Scatter plot of significantly regulated genes in soleus compared to GC. (B) GO enrichment analysis of pathways in GC. (C) Relative expression levels of Trim63 and Fbxo32. (D-G) Metabolomics of GC from *Nfe2l1* mKO and WT mice (*n* = 10 - 14 mice per group). (G) PCA of metabolic phenotypes. (H) Heatmap of the metabolome profile. (I) Top enriched metabolite sets in GC muscle tissue. RNA-seq data are mean ±SEM, $P < 0.01$ by *t*-test (A-C) and metabolomics data are mean ±SEM, $P < 0.05$ by *t*-test (D-G).

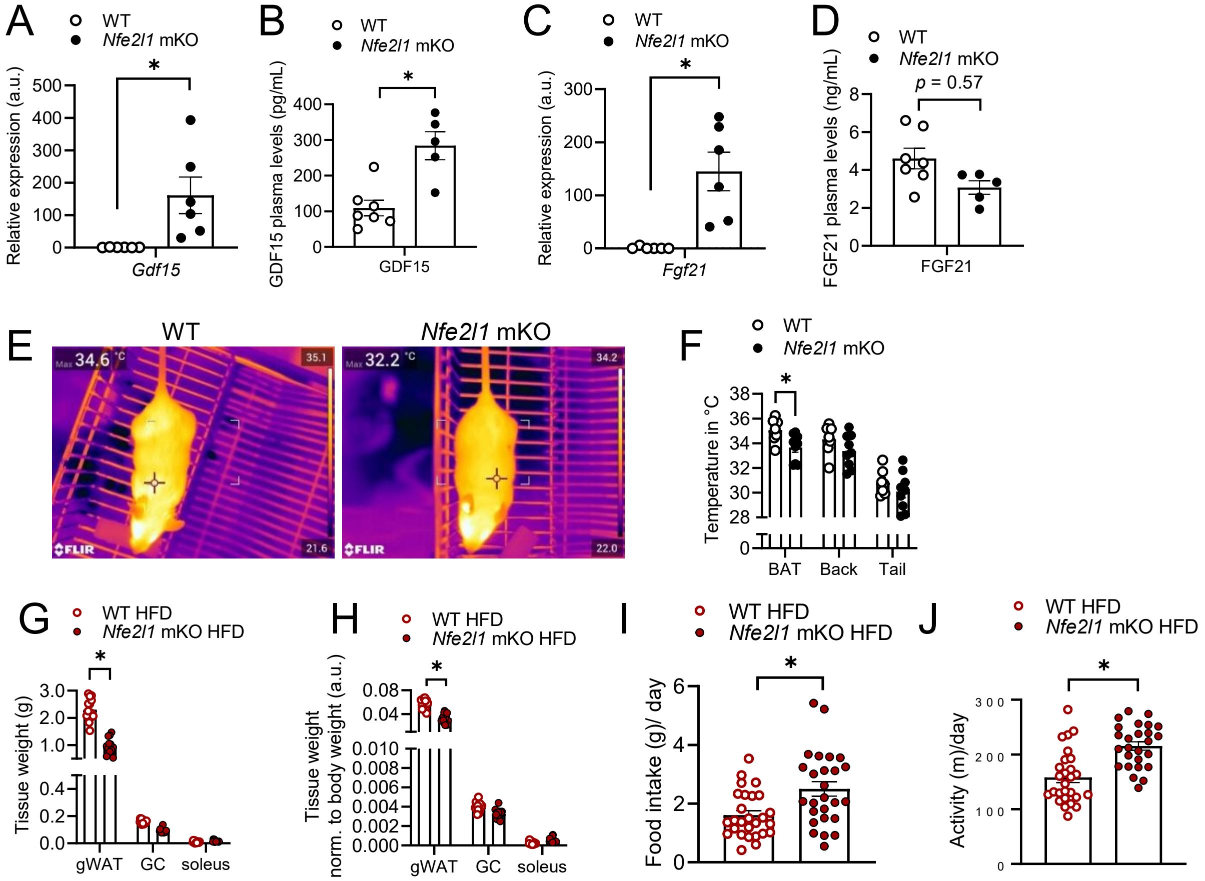


**Figure EV4: Myocyte Nfe2l1 is a critical regulator of energy expenditure and food intake.**

(A) Relative *Gdf15* mRNA expression and (B) plasma levels of GDF15 in WT and mKO mice. (C) Relative *Fgf21* mRNA expression and (D) plasma levels of FGF21 in *Nfe2l1* mKO mice and WT controls. ((A-D) *n* = 5 -7 mice per group). (E) Representative forward-looking infrared (FLIR) images and (F) Corresponding measurements of temperature around interscapular BAT depot, back and base of the tail at 3 months of age from *Nfe2l1* mKO animals and WT littermates (*n* = 7-9 animals per group). (G-J) Assessment of *Nfe2l1* mKO and WTs after 16 weeks of HFD. (G) Tissue weight of gWAT, GC and soleus and (H) Tissue weight normalized to body weight of mKO and WTs on HFD (*n* = 14 – 15 mice per group). (I) Food intake and (J) activity (*n* = 26 – 27 mice per group). Data are mean ±SEM, $P < 0.05$ by *t*-test (A-D, I, J) or 2-way ANOVA (F-H).

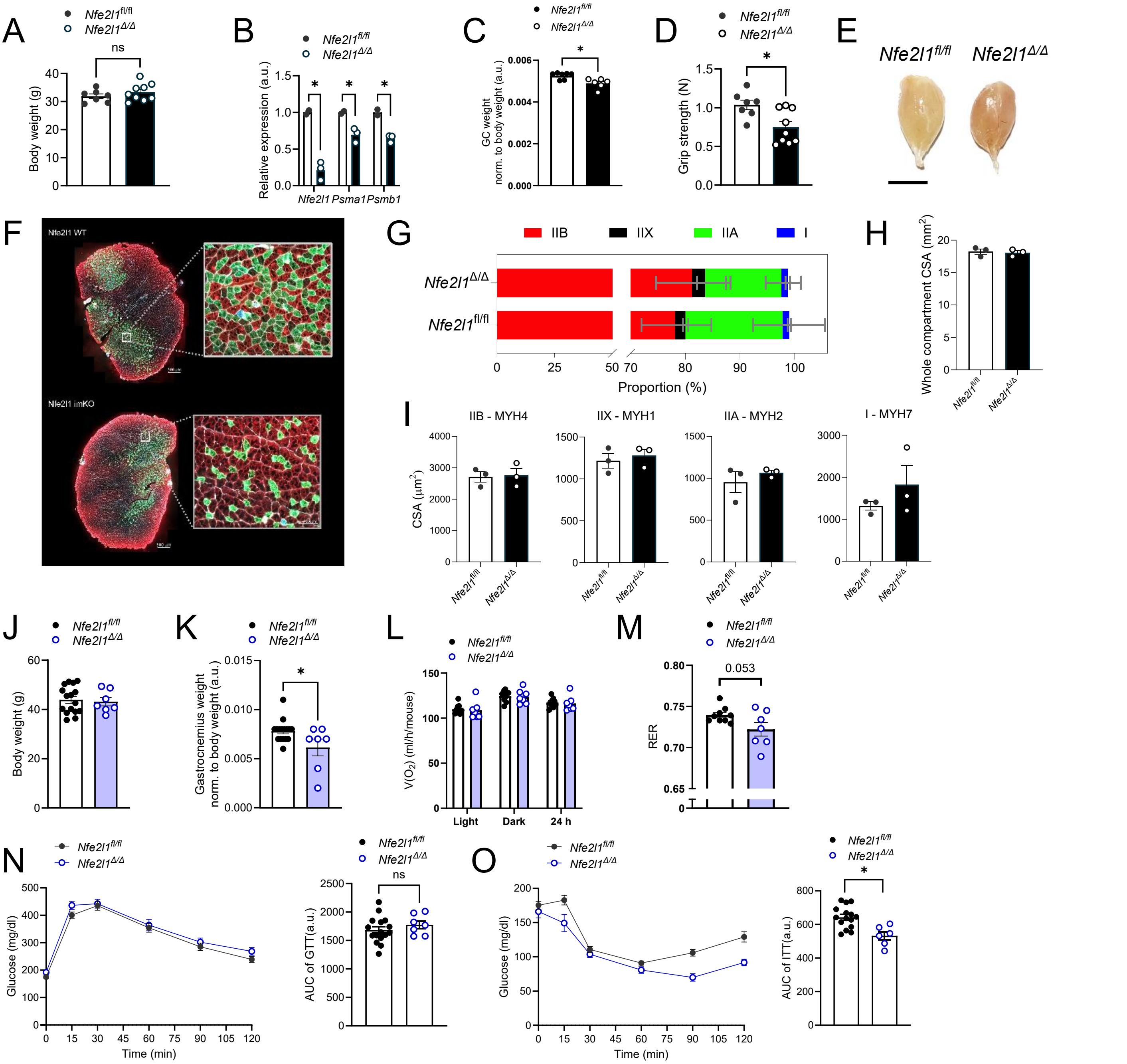


**Figure EV5: Absence of myocyte Nfe2l1 in adult mice impacts muscle phenotype and improves insulin sensitivity.**

(A-I) Assessment of the tamoxifen inducible Nfe2l1 Acta1-CreERT2 model, with Cre negative (Nfe2l1$^{fl/fl}$) and the Cre positive (Nfe2l1$^{\Delta/\Delta}$) mice, 8 weeks after injection. (A) Body weight ($n$ = 7 - 9 mice per group). (B) Relative expression levels of Nfe2l1 and downstream targets Psma1 and Psmb1 ($n$ = 3 mice per group). (C) Tissue weight of the Gastrocnemius muscle (GC) normalized to body weight ($n$ = 6 - 7 mice per group). (D) Forelimb grip strength ($n$ = 7 - 9 mice per group). (E) Representative image of GC muscle. Scale: 0.5 cm. (F) Representative immunofluorescence images of fiber types stained with MyHC-2B (red), MyHC-2A (green), MyHC-1 (blue) in GC from WT controls and Nfe2l1$^{\Delta/\Delta}$ mice (G) Proportion of fibers in GC muscle. (H) Assessment of whole compartment and (I) Fiber type specific cross-sectional area (CSA) from fluorescent imaging. (J-O) Analysis of tamoxifen inducible Nfe2l1 Acta1-CreERT2 model, with Cre negative (Nfe2l1$^{fl/fl}$) and the Cre positive (Nfe2l1$^{\Delta/\Delta}$) mice after tamoxifen injection and consequent HFD feeding for 16 weeks. (J) Body weight of WT controls and Nfe2l1$^{\Delta/\Delta}$ mice. (K) Gastrocnemius weight normalized to body weight. (L) Oxygen consumption and (M) Respiratory exchange ratio (RER) measured by indirect calorimetry. (N) Traces and quantification (area under curve (AUC)) of intraperitoneal glucose tolerance test (GTT). (O) Traces and quantification (AUC) of intraperitoneal insulin tolerance test (ITT). (J-O: ($n$ = 7 - 16 mice per group)). The data are mean ±SEM, $p<0.05$ by two-way ANOVA (B) or $t$-test (A,C,D,I-O ).
