## Supplementary figures and images for "The myocyte Nfe2l1-ubiquitin-proteasome system controls muscle fiber type and obesity-induced insulin resistance"

### Supplemental material (uncropped blots)

#### Uncropped WBs

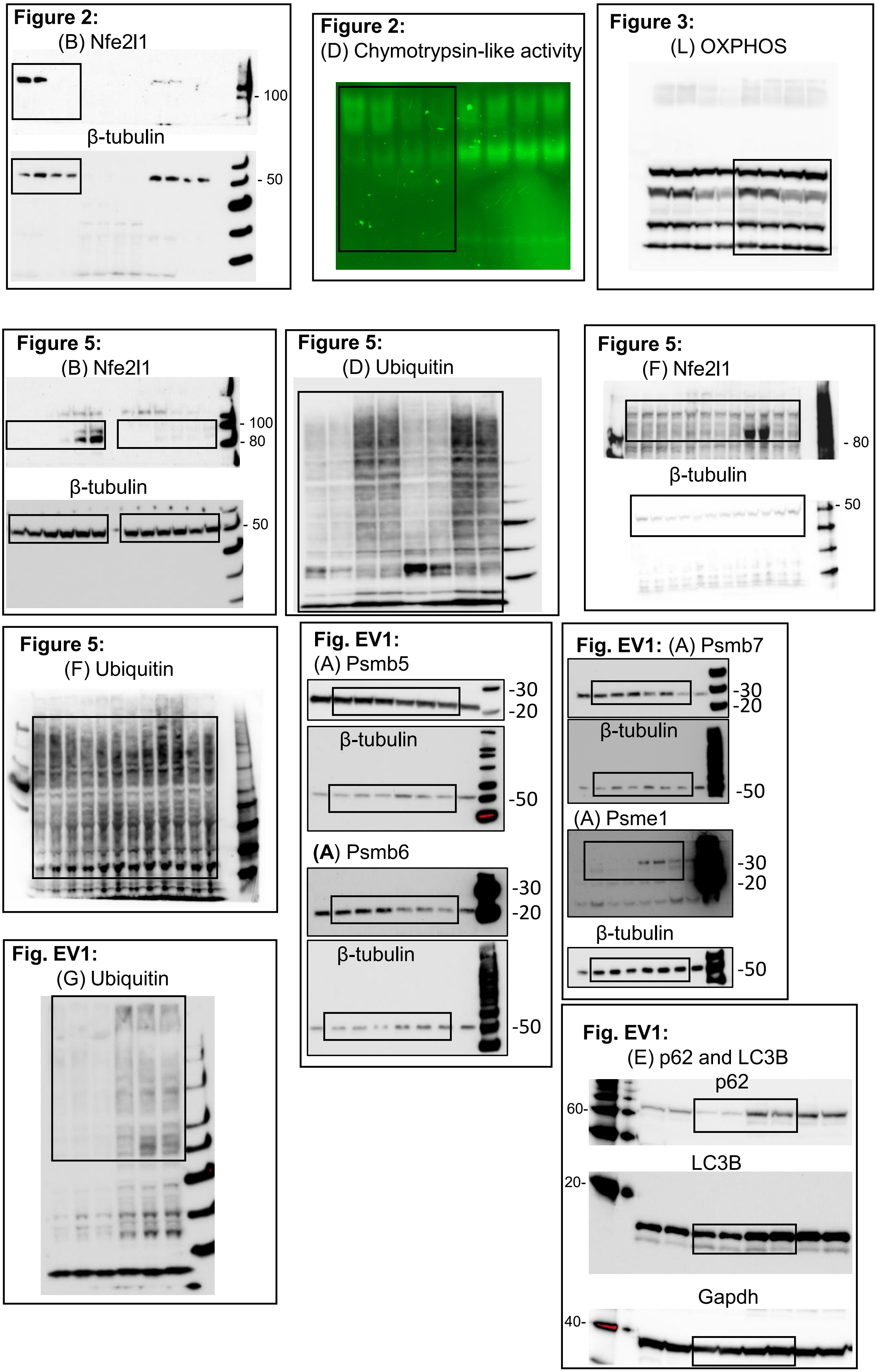
